# Cognitive and Synaptic Impairment Induced by Deficiency of Autism Risk Gene *Smarcc2* and its Rescue by Histone Deacetylase Inhibition

**DOI:** 10.1101/2025.05.29.656867

**Authors:** Pei Li, Siqi Men, Prachetas J. Patel, Komal Saleem, Ping Zhong, Kin Wai Tam, Jian Feng, Zhen Yan

**Affiliations:** Department of Physiology and Biophysics, Jacobs School of Medicine and Biomedical Sciences, State University of New York at Buffalo, Buffalo, NY 14203, USA

**Keywords:** autism, Smarcc2, cognitive function, synaptic transmission, GABA, glutamate, histone acetylation, epigenetics

## Abstract

*SMARCC2*, which encodes BAF170, a core subunit of chromatin remodeling BAF complex, is one of the top-ranking risk genes for autism spectrum disorder (ASD). However, the mechanisms linking *SMARCC2* haploinsufficiency to ASD remain poorly understood. Genome-wide RNA-seq analysis revealed that *SMARCC2* was significantly diminished in iPSC-derived neurons from idiopathic ASD patients. ChIP-seq of *SMARCC2* demonstrated its binding to many other ASD risk genes involved in transcriptional regulation. *Smarcc2* deficiency in prefrontal cortex (PFC) of adolescent mice led to impaired working memory, with largely intact social and anxiety-like behaviors. Significant downregulation of genes enriched in synaptic transmission were found in PFC of S*marcc2*-deficient mice by RNA-seq and qPCR profiling. In parallel, electrophysiological recordings uncovered the significant impairment of GABAergic and glutamatergic synaptic currents in S*marcc2*-deficient PFC pyramidal neurons. Smarcc2 bound to HDAC2, and *Smarcc2* deficiency led to the reduced global histone acetylation and H3K9ac enrichment at synaptic gene *Slc1a3* (EAAT1), *Slc6a1* (GAT1), and *Slc32a1* (VGAT) promoters. Treatment of S*marcc2*-deficient mice with romidepsin, a class I HDAC inhibitor, restored histone acetylation, working memory and some synaptic gene expression. These findings highlight the critical role of Smarcc2 in regulating cognitive and synaptic function, suggesting that targeting HDAC could alleviate deficits in Smarcc2-associated neurodevelopmental disorders.

## Introduction

Autism Spectrum Disorder (ASD) and intellectual disability (ID) are complex neurodevelopmental disorders, with 30%-80% ASD cases accompanied by ID [1]. Growing evidence highlights the significant contribution of genetic factors to both ASD and ID [2–4]. Many of the top-ranking ASD risk genes function as histone modifiers and chromatin remodelers [2], suggesting the critical role of epigenetic mechanisms in ASD pathogenesis and therapeutic intervention [5–10].

*SMARCC2*, which encodes BAF170, a key subunit of the ATP-dependent chromatin remodeling BAF complex (mammalian SWI/SNF complex) [11, 12], is one of the top-ranking ASD risk genes [2, 13]. The SWI/SNF subfamily provides nucleosome rearrangement, enabling binding of specific transcription factors [14] and allowing genes to be activated or repressed [15]. Haploinsufficiency of Smarcc2 causes a syndrome characterized by intellectual disability, developmental delay, and ASD [16, 17]. Preclinical studies suggest that Smarcc2 plays a critical role in early development of forebrain [18], as well as proliferation and differentiation of neural stem cells in dentate gyrus of hippocampus [19].

It remains unknown whether Smarcc2 deficiency is linked to synaptic dysfunction underlying ASD/ID phenotypes. Thus, we used the virus-mediated approach to knock down Smarcc2 in adolescent mice and examined the consequences at genomic, electrophysiological and behavioral levels. Prefrontal cortex (PFC), which controls cognitive & emotional processes and is highly disrupted in ASD [20–22], was selected for Smarcc2 knockdown. In addition, we uncovered a therapeutic strategy to alleviate the abnormalities of Smarcc2-deficient mice.

## Results

### *SMARCC2* is downregulated in iPSC-derived neurons from ASD patients, and *SMARCC2*- binding genes are enriched in ASD risk factors

To find out gene alterations in ASD, we generated iPSC-derived cortical neurons from idopathic ASD patients and control subjects, and performed RNA-seq to examine genome-wide changes. As shown in **Fig. 1a**, our iPSC-derived cortical neurons had elaborate dendritic processes and formed well-connected networks with the expression of neuronal markers MAP2 and TUJ1. RNA-seq revealed a number of differentially expressed genes (DEGs) in ASD iPSC-derived neurons (668 downregulated, 906 upregulated, **Fig. 1b, Sup. Table 1**). The downregulated genes were enriched in transcriptional regulation and chromatin organization (**Fig. 1c**), including many top-ranking ASD risk genes, such as *CHD8, SMARCC2, CHD2, KDM6B, ASH1L, CREBBP, POGZ,* and *FOXP1* (**Fig. 1d**). As shown in **Fig. 1e**, *SMARCC2* expression was significantly diminished in iPSC-derived neurons from ASD patients.

**Figure 1.**
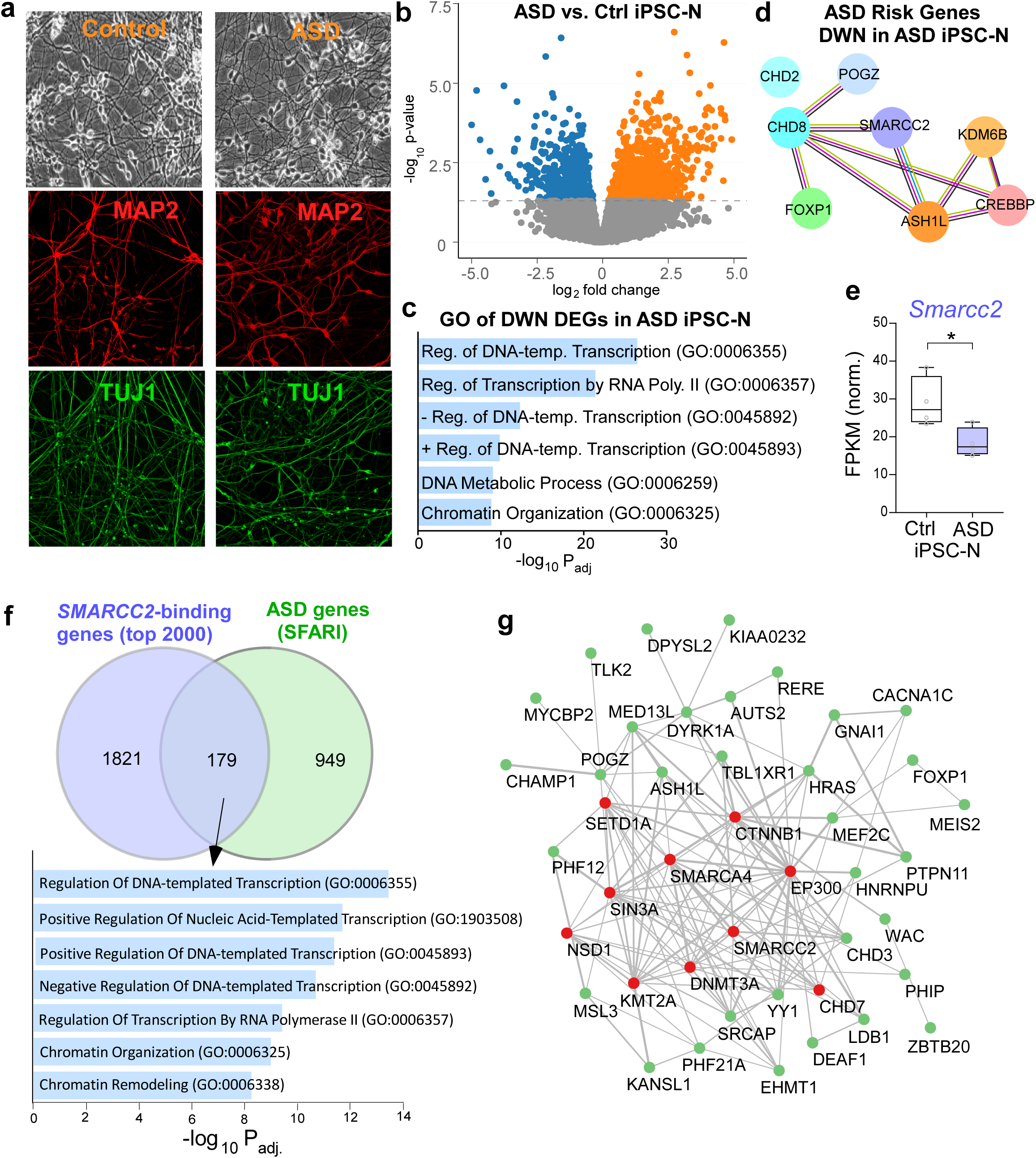
*SMARCC2* is downregulated in iPSC-derived neurons from ASD patients, and it binds to other ASD risk genes. **(a)** Phase contrast and immunocytochemical images of iPSC-derived neurons from a control and an ASD patient (Day 60) stained with neuronal markers (MAP2 and TUJ1). **(b)** Volcano plot showing differentially expressed genes (DEGs) in iPSC-derived neurons from ASD vs. control subjects (n=4 lines/group). **(c)** Gene Ontology (GO) enriched pathways of downregulated genes in iPSC-derived neurons from ASD patients. **(d)** Protein-protein interaction (PPI) network of top-ranking ASD risk genes downregulated in iPSC-derived neurons from ASD patients. **(e)** Box plots of *SMARCC2* mRNA levels in iPSC-derived neurons from control vs. ASD patients (*p*=0.016, n=4 lines/group, t-test). \**p*<0.05. **(f)** Venn diagrams showing the significant overlap of top 2000 *SMARCC2*-binding genes identified from ChIP-seq data with ASD risk genes from SFARI database. Inset: Bar graphs of GO pathways enrichment for overlapping genes. **(g)** PPI network of *SMARCC2* binding genes that belong to category 1 SFARI ASD risk genes.

Among the top 2000 genes with *SMARCC2* binding at their promoters (1 kb of TSS), we found that 179 genes overlapped with SFARI ASD gene dataset (1128 genes) (1.59 fold over-enrichment, hypergeometric p-value: 1.64E-10, **Sup. Table 2**). Gene Ontology (GO) enrichment analysis revealed that these overlapping genes had significant enrichment in pathways related to DNA-templated transcription regulation, gene expression, and chromatin remodeling (**Fig. 1f**). Among the 232 SFARI category I ASD genes, 43 had *SMARCC2* binding at their promoters, and they formed a well-connected protein-protein interaction (PPI) network (**Fig. 1g**). Hub genes in the PPI network included histone acetyltransferase (*EP300*), histone methyltransferases (*SETD1A*, *KMT2A*, *NSD1*), chromatin remodelers (*SMARCA4, CHD7*), DNA methyltransferase (*DNMT3A*), transcriptional repressor (*SIN3A),* and dual functional gene involved in cell-cell adhesion and transcription (*CTNNB1*). These findings suggest that SMARCC2 could regulate gene expression and cellular function directly or indirectly through other interacting ASD genes.

### *Smarcc2* deficiency in PFC impairs cognitive function without affecting other behaviors

To find out how *SMARCC2* haploinsufficiency is linked to ASD, we performed the virus-based knockdown of Smarcc2. Smarcc2 mRNA was significantly reduced in N2A cells transfected with *Smarcc2* shRNA, confirming the knockdown (KD) effectiveness (**Fig. 2a**). Next, *Smarcc2* shRNA AAV was stereotaxically injected bilaterally into the medial PFC of 5-week-old C57BL/6 mice (**Fig. 2b**). Two weeks post-injection, we detected the significant reduction of Smarcc2 mRNA and protein levels in the PFC of Smarcc2^KD^ mice (**Fig. 2b**). A marked decrease of Smarcc2 fluorescence intensity was also observed in immunostaining, while the number of NeuN+ neurons was unchanged by Smarcc2 knockdown (**Fig. 2c** and **2d**).

**Figure 2.**
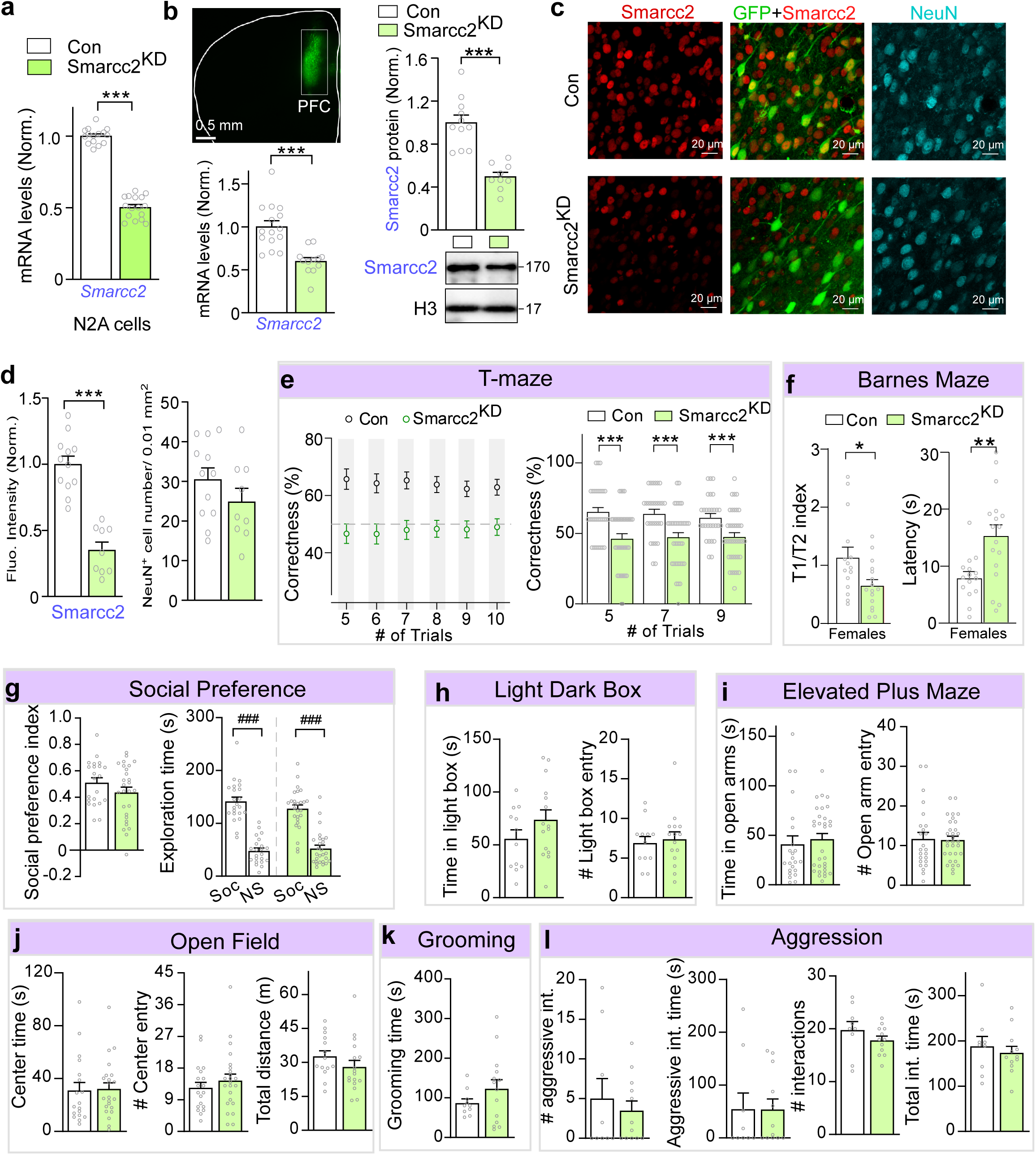
Smarcc2 deficiency in PFC induces cognitive deficits without affecting other behaviors. **(a)** Bar graphs showing Smarcc2 mRNA levels in N2A cells transfected with control vs. Smarcc2 shRNA. n=15-16 cultures/group; t_29_=20.3, *p*<0.001, unpaired t test. **(b)** Representative low-magnification (2×) image showing the AAV injection site and bar graphs displaying Smarcc2 mRNA and protein levels in mouse PFC injected with control vs. Smarcc2 shRNA AAV. Inset: immunoblots of Smarcc2 and Histone H3. mRNA: n=12-15 mice/group; t_25_=4.78, *p*<0.001. Protein: n=9-11 mice/group; t_18_=5.9, *p*<0.001. Scale bars: 0.5 mm. **(c)** Representative confocal images (63×) showing immunostaining of Smarcc2 (red) and NeuN (blue). Scale bars: 20 μm. **(d)** Quantification of the fluorescence intensity of Samrcc2 and the number of NeuN+ cells. n=9-12 images/3-4 mice/ group. Smarcc2 intensity: t_19_=7.4, *p*<0.001, unpaired t test. **(e)** Plot showing the percentage of correct responses in various trials of spontaneous T-maze tests of control vs. Smarcc2^KD^ mice. n=31-41 mice/group. *p*<0.001, unpaired t test; **(f)** Bar graphs showing the spatial memory index and latency to the correct hole in Barnes Maze (BM) tests of control vs. Smarcc2^KD^ female mice. n=15-16 mice/group. Index: *p*=0.03, M-W test; latency: t_23.6_=3.26, *p*=0.034, Welch’s t test. **(g)** Bar graphs showing the social preference index and time spent on social (Soc) and non-social (NS) objects in three-chamber social preference tests of control vs. Smarcc2^KD^ mice. n=22-27 mice/group. Time: F_1,94_=147.5, ^###^*p*<0.001, Soc vs. NS, two-way ANOVA. **(h)** Bar graphs showing the time spent in light box and the number of entries into the light box in light-dark box tests of control vs. Smarcc2^KD^ mice. n=12-15 mice/group. **(i)** Bar graphs showing the time in open arms and number of entries into open arms in elevated plus maze (EPM) tests of control vs. Smarcc2^KD^ mice. n=24-31 mice/group. **(j)** Bar graphs showing time spent in the center area of open field, the number of entries to center area, and the total distance traveled in open field (OF) tests of control vs. Smarcc2^KD^ mice. n=18-22 mice/group. **(k)** Bar graph showing the grooming time in grooming tests of control vs. Smarcc2^KD^ mice. n=10-14 mice/group. **(l)** Bar graphs showing the number and time of aggressive encounters and total interactions in resident-intruder (RI) tests of control vs. Smarcc2^KD^ mice. n=9-12 mice/group. All data are shown as mean ± SEM, \**p*<0.05, \*\*\**p*<0.001.

Next, we assessed behavioral consequences of Smarcc2 knockdown. Since *de novo SMARCC2* variants are associated with ID, we first evaluated cognitive function by examining working memory using spontaneous T-maze alternation tests and spatial memory using Barnes Maze (BM) tests. Compared to control mice, correctness in T-maze tests of Smarcc2^KD^ mice were significant decreased across trials (**Fig. 2e**), with both male and female Smarcc2^KD^ mice displaying impaired working memory (**Fig. S1a**). In BM tests, female Smarcc2^KD^ mice exhibited a significant reduction of spatial memory index (Time on correct hole / Time on all incorrect holes) and an increased latency to the correct hole (**Fig. 2f**), whereas male Smarcc2^KD^ mice showed no significant changes (**Fig. S1b**), suggesting that only female Smarcc2^KD^ mice exhibited spatial memory deficits.

Given the strong association of Smarcc2 haploinsufficiency to autism, we further assessed social behavior using three-chamber social preference tests. No significant alteration was found in Smarcc2^KD^ mice on social preference index and the time spent interacting with a social or non-social objects (**Fig. 2g**), albeit a slight reduction of social preference index in female Smarcc2^KD^ mice (**Fig. S1c**), suggesting that sociability is largely normal.

We also examined anxiety-related behaviors using light-dark box (LDB) tests, elevated plus maze (EPM) tests, and open field (OF) tests. In all these measurements, compared to controls, Smarcc2^KD^ mice displayed no significant differences (**Fig. 2h-2j**) in both sexes (**Fig. S1 d-f**).

Additionally, self-grooming, a marker of repetitive behavior, was unaffected by Smarcc2 KD (**Fig. 2k**). In resident-intruder tests, male Smarcc2^KD^ mice exhibited no change in the frequency or time of aggressive encounters or total interactions (**Fig. 2l**).

### *Smarcc2* deficiency induces the downregulation of synaptic genes

Since Smarcc2 is a core subunit of BAF chromatin remodeler complex crucial for gene regulation [23], we performed RNA sequencing (RNA-seq) analysis to assess genome-wide transcriptional changes following Smarcc2 knockdown. Using the cutoff of fold change ≥ 20% and p value ≤ 0.05, we identified 1735 differentially expressed genes (DEGs), with 653 downregulated and 1082 upregulated genes in Smarcc2^KD^ mice (**Fig. 3a, Sup. Table 3**). GO analyses revealed that these downregulated genes were primarily enriched in synaptic transmission, vascular transport, and signal release from synapse (**Fig. 3b**). Synaptic enrichment analysis with SynGO, a synaptic gene ontology database, identified presynaptic and postsynaptic genes as the most enriched components downregulated after Smarcc2 KD (**Fig. 3c**). PPI of downregulated synaptic genes by Smarcc2 KD (**Fig. 3d**) can be classified into 5 major categories, (1) glutamatergic synapse genes, including AMPA receptor (*Gria3*), NMDA receptor (*Grin2b*), glutamate transporters (*Slc1a2*, *Slc1a3*, and *Slc38a3*), (2) GABAergic synapse genes, including GABA receptors and transporters (*Gabbr1*, *Gabrb3*, *Slc6a11*, *Slc32a1*, and *Slc38a3*), GABA-synthesizing enzyme (*Gad1*), (3) synaptic vesicle release genes, including *Nrxn1*, *Syt1*, *Syn1*, *Cplx1*, *Cplx3*, and *Dnajc5*, (4) ion channels, including sodium channels (*Scn1b*, *Scn4b*, and *Scn8a*), nicotinic acetylcholine receptor (*Chrnb2*), calcium channel (*Cacng2*), potassium channel (*Kcnq3*), (5) monoamine receptors, including serotonin receptors (*Htr1b*, *Htr5a*, and *Htr3a*), and dopamine receptors (*Drd1* and *Drd2*).

**Figure 3.**
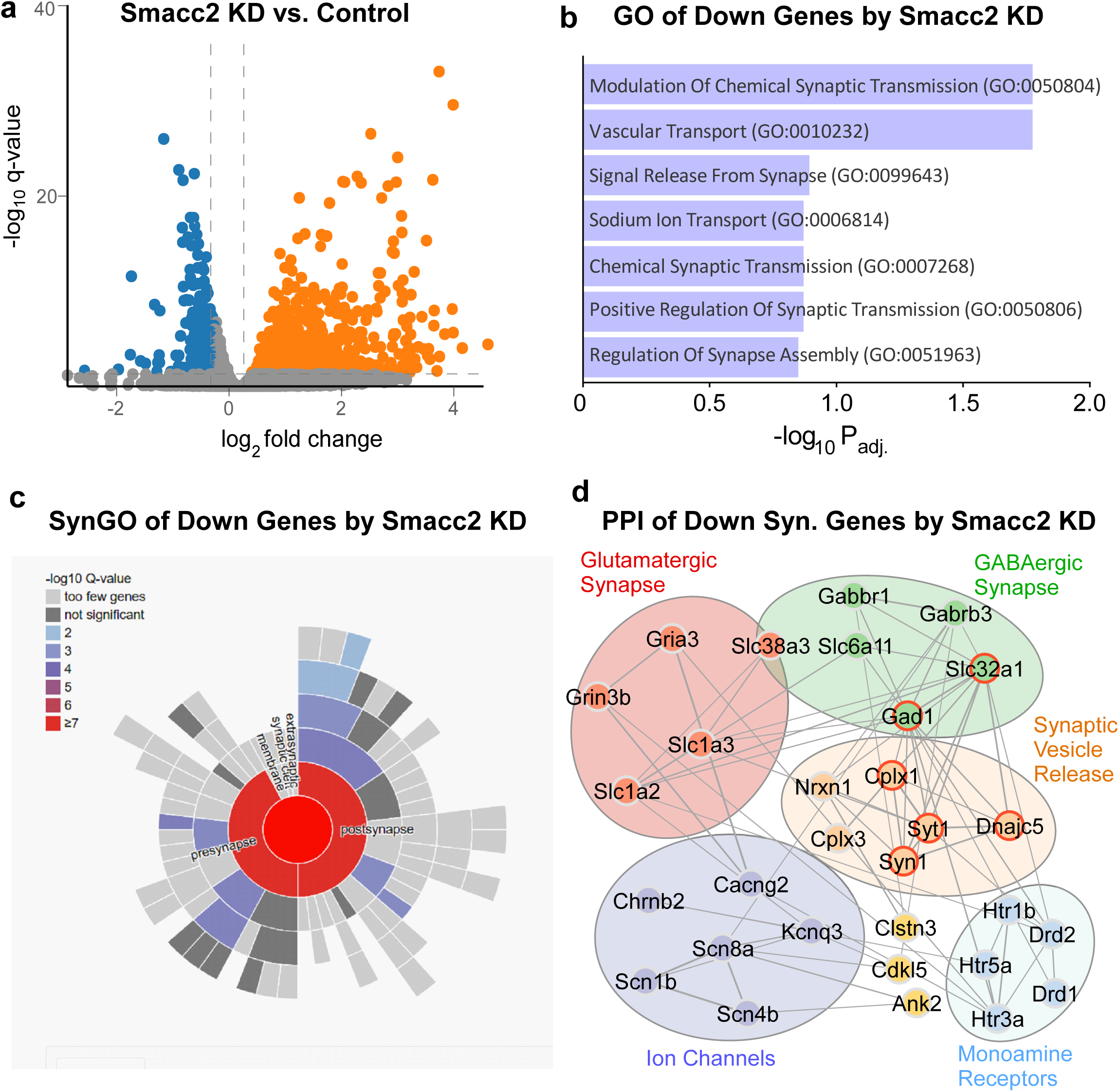
RNA-seq reveals that Smarcc2 deficiency downregulates synaptic genes in PFC. **(a)** Volcano plot showing differentially expressed genes (DEGs) in PFC of Smarcc2^KD^ mice, compared to controls. **(b)** Gene Ontology (GO) enriched pathways of downregulated genes by Smarcc2 KD. **(c)** Sunburst plot from SynGO analysis illustrating the most enriched cellular component categories among the downregulated genes in top GO categories in Smarcc2^KD^ mice. Higher red intensities indicate greater significance. **(d)** Protein-protein interaction (PPI) network of downregulated synaptic genes in Smarcc2^KD^ mice. Hub proteins are circled in red.

Our analysis of the RNA-seq data of embryonic cells from Smarcc1/2 double knockout mice [18] also found that the significantly downregulated genes were enriched in chemical synaptic transmission following nervous system development and axon guidance (**Fig. S2a** and **S2b, Sup. Table 4**). They formed a well-connected PPI network including glutamatergic synapse genes, GABAergic synapse genes, and presynaptic genes (**Fig. S2c**), some of which overlapped with our current study (e.g. *Slc1a2/3*, *Gria3*, *Nrxn1*, *Gabrb3*, *Cplx1*).

Notably, Smarcc2 knockdown also induced the upregulation of genes, which were enriched in extracellular matrix organization, immune response, phagocytosis, and apoptosis (**Fig. S3a**). PPI of upregulated genes in top GO pathway pointed to key collagen genes (*Col1a1*, *Col15a1*, *Col18a1, etc.*) and matrix metalloproteinases (*Mmp14, Mmp2*) (**Fig. S3b**).

Guided by RNAseq data and bioinformatic analysis, we used qPCR to examine more synaptic genes in larger sized samples. For GABA-related genes, a significant decrease was found in Smarcc2^KD^ mice on *Slc6a1* (encoding GAT1), *Slc32a1* (encoding VGAT), and *Gad1*, while GABA receptor genes (*Gabra4*, *Gabrb2*, *Gabrg2*, *Gabrg3)*, *Bsn* (encoding Bassoon), *Sst* (encoding Somatostatin), *Pvalb* (encoding Parvalbumin) and *Vip* (encoding VIP) were largely unchanged (**Fig. 4a**). For glutamate-related genes, a significant decrease was found in Smarcc2^KD^ mice on *Gria3* (encoding AMPAR subunit GluR3) and *Slc1a3* (encoding EAAT1), while other glutamate receptor subunits (*Gria1*, *Gria2*, *Grin1*, *Grin2a*, *Grin2b, Grm2),* transporter or synaptic organizers (*Slc17a6*, *Homer1a*, *Nrxn1*, *Dlgap4*) were not significantly changed. Presynaptic genes for synaptic vesicle trafficking and exocytosis (*Snap25*, *Stxbp1*, *Stxbp5*, *Syp*) were also not significantly changed (**Fig. 4b**). These data suggest that Smarcc2 deficiency in PFC of juvenile mice induces the robust reduction of only a subset of synaptic genes.

**Figure 4.**
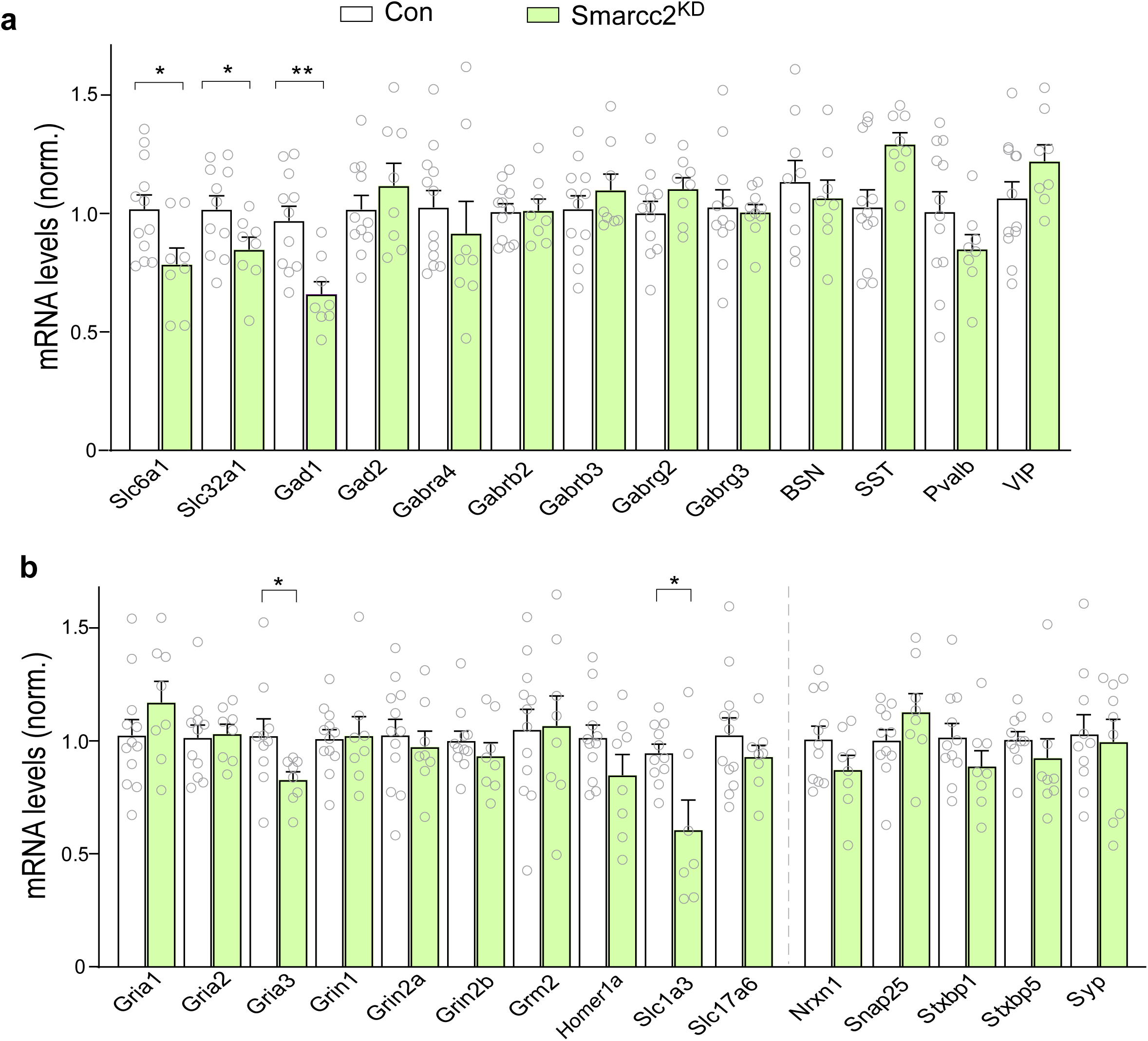
Quantitative PCR profiling confirms the downregulation of some synaptic genes in PFC of Smarcc2^KD^ mice. **(a, b)** Bar graphs showing the mRNA level of synaptic genes (GABAergic, glutamatergic and presynaptic) in PFC of control vs. Smarcc2^KD^ mice. n=8-12 mice/group. All data are shown as mean ± SEM, \**p*<0.05, \*\**p*<0.01.

### *Smarcc2* deficiency in PFC diminishes inhibitory and excitatory synaptic transmission

Given the downregulation of synaptic genes in Smarcc2^KD^ mice, we next performed whole-cell parch-clamp recordings of layer V pyramidal neurons in PFC slices to examine the physiological impact of Smarcc2 deficiency. As shown in **Fig. 5a**, Smarcc2^KD^ mice had a significant decrease of the amplitude and frequency of spontaneous inhibitory postsynaptic current (sIPSC), compared to control mice. Furthermore, Smarcc2^KD^ mice exhibited a marked reduction in the input/output curve of IPSC evoked by a series of stimulation intensities (**Fig. 5b**). Paired-pulse ratio (PPR) of evoked IPSC, a measure of presynaptic GABA release, was not significantly changed (**Fig. 5c**). In parallel experiments examining excitatory transmission, Smarcc2^KD^ mice had a significant decrease of spontaneous excitatory postsynaptic current (sEPSC) amplitude and frequency (**Fig. 5d**). Also, Smarcc2^KD^ mice exhibited the decreased EPSC evoked by various stimuli (**Fig. 5e**), while PPR of evoked EPSC was largely unchanged (**Fig. 5f**). These data suggest that GABAergic synaptic inhibition and glutamatergic synaptic excitation are compromised by SMARCC2 deficiency, probably due to the attenuated expression of synaptic genes.

**Figure 5.**
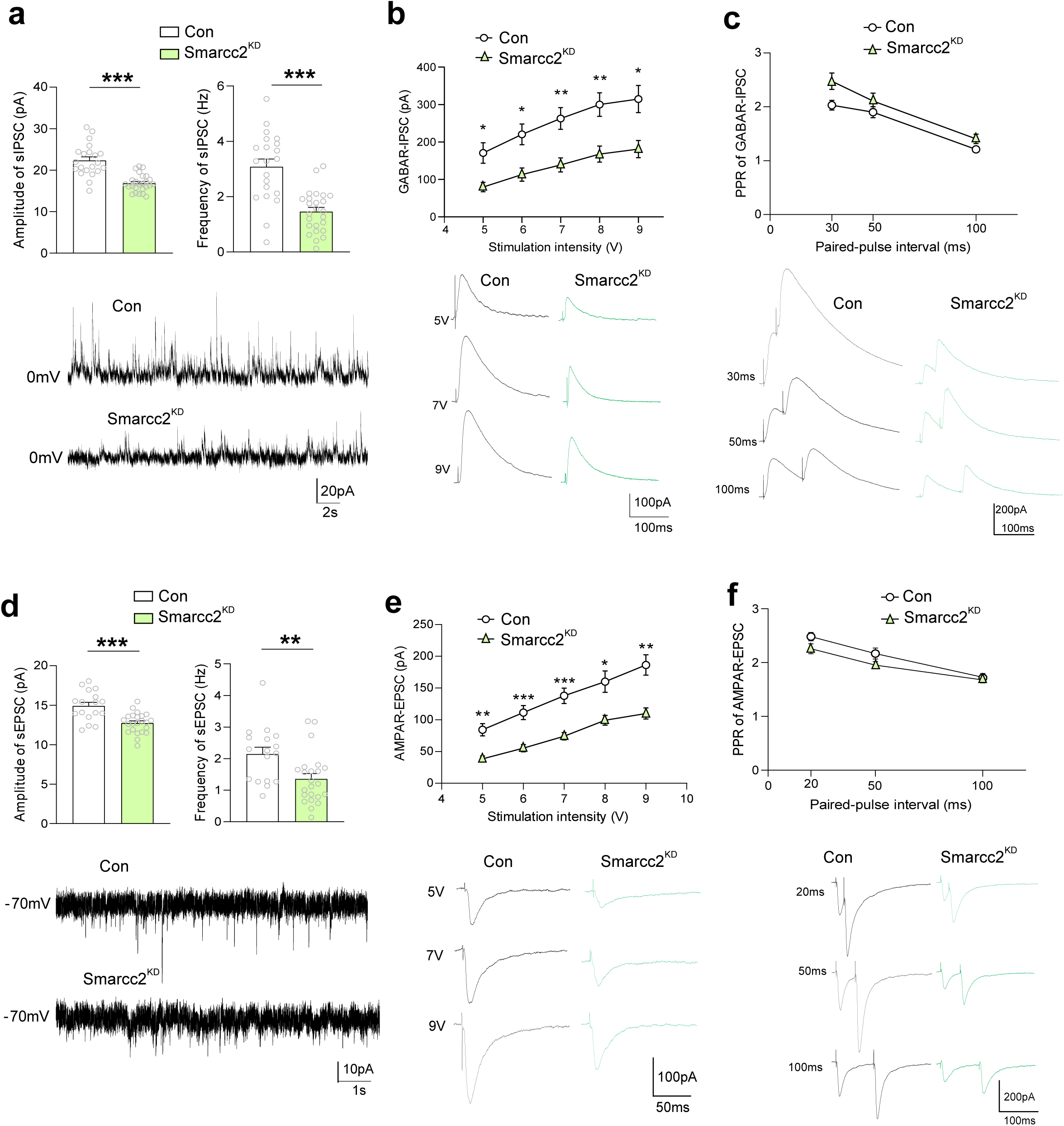
Smarcc2 deficiency impairs GABAergic and Glutamatergic transmission of PFC pyramidal neurons. **(a-c)** Plots of spontaneous IPSC (sIPSC) amplitude and frequency (a), input-output curves of evoked IPSC (b), and paired-pulse ratio (PPR) of IPSC (c) in PFC pyramidal neurons from control vs. Smarcc2^KD^ mice. a, t_29.3_ = 5.9 (amp), t_31.3_ = 5.2 (freq), *p*<0.001, two-tailed t test, n=21-25 cells/ 5-6 mice/ group; b, F_1,31(genotype)_ = 12.5, p=0.0013, *p*<0.05, *p*<0.01, two-way rmANOVA, n=16-17 cells/ 5 mice/ group; c, F_1,29(genotype)_ = 5.0, p=0.033, two-way rmANOVA, n=15-16 cells/ 5 mice/ group. Inset: representative sIPSC and eIPSC traces. **(d-f)** Plots of spontaneous EPSC (sEPSC) amplitude and frequency (d), input-output curves of evoked EPSC (e), and PPR of EPSC (f) in PFC pyramidal neurons from control vs. Smarcc2^KD^ mice. d, t_38_ = 4.3 (amp), *p*<0.01; t_38_ = 2.9 (freq), *p*<0.001, two-tailed t test, n=17-23 cells/5-6 mice/group; e, F_1,29(genotype)_ = 20.9, *p*<0.001, two-way rmANOVA, n=15-16 cells/5 mice/group; f, F_1,35(genotype)_ = 3.0, p=0.09, two-way rmANOVA, n=18-19 cells/5 mice/group. Inset: representative sEPSC and eEPSC traces. Data are shown as mean ± SEM, \**p*<0.05, \*\**p*<0.01, \*\*\**p*<0.001.

### *Smarcc2* deficiency induces the reduction of histone acetylation

Among Smarcc2-binding genes involved in ASD, many are histone modifiers, including EP300, ASH1L, EHMT1, KMT2A, SETD1A (Fig. 1b). Thus, we tested whether Smarcc2 deficiency might affect histone acetylation and methylation, therefore altering gene expression. In N2A cells transfected with Smarcc2 shRNA, we found a significant decrease of the level of H3K9ac and H3K27ac, while H3K27me3 and H3K4me3 levels were unchanged (**Fig. 6a**). Similarly, significantly decreased H3K9ac and H3K27ac levels were found in PFC of mice injected with Smarcc2 shRNA AAV, without significant changes in H3K27me3 and H3K4me3 levels (**Fig. 6b**).

**Figure 6.**
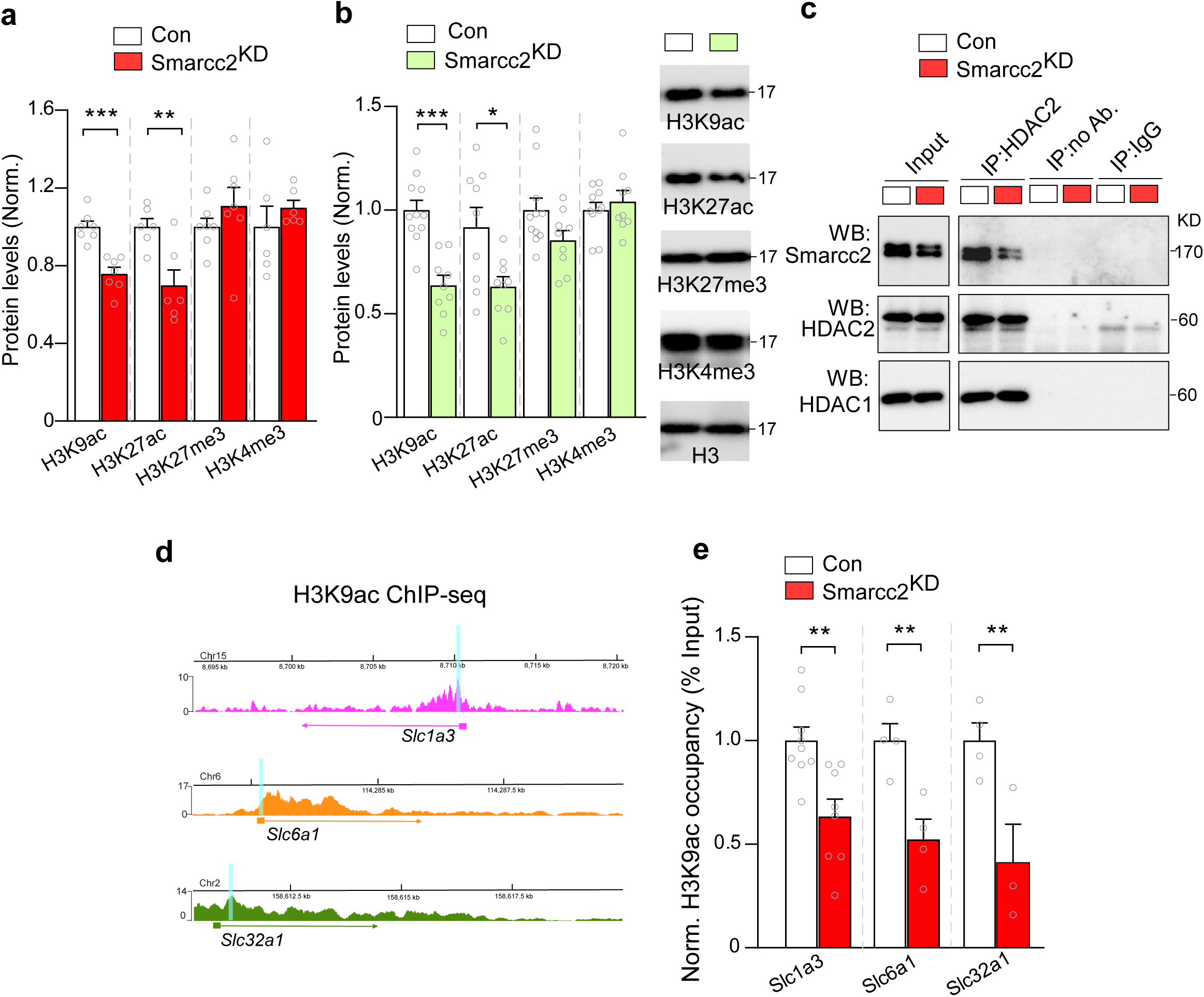
Smarcc2 deficiency reduces global histone acetylation, diminishes HDAC2-Smarcc2 binding, and decreases H3K9ac occupancy at synaptic gene promotors. **(a, b)** Bar graphs showing the level of H3K9ac, H3K27ac, H3K27me3, and H3K4me3 in N2A cells transfected with control vs. Smarcc2 shRNA (a) or PFC tissue from mice injected with control vs. Smarcc2 shRNA AAV (b). a, n=6-7 cultures/group, H3K9ac: t_12_=5.5, *p*<0.001; H3K27ac: t_10_=3.3, *p*=0.008; b, n=9-11 mice/group, H3K9ac: t_18_=5.3, *p*<0.001; H3K27ac: t_16_=2.7, *p*=0.016. (c) Representative immunoblots of co-IP experiments showing HDAC2-bound Smarcc2 in control vs. Smarcc2^KD^ mice. Anti-HDAC2 was used for IP, anti-Smarcc2, anti-HDAC2 and anti-HDAC1 was used for blotting. No antibody and IgG were used as negative controls for IP. **(d)** Diagram showing H3K9ac occupancy at *Slc1a3, Slc6a1,* and *Slc32a1* promotor regions, including the location of primers used in ChIP-PCR assays (light blue rectangular area). **(e)** ChIP assay data showing differential H3K9ac levels at *Slc1a3, Slc6a1* and *Slc32a1* promoters in the PFC of control vs. Smarcc2^KD^ mice. n=3-6 mice/group. *Slc1a3*: t_15_=3.5, *p*=0.003; *Slc6a1*: t_6_=3.8, *p*=0.009; *Slc32a1*: t_5_=3.2, *p*=0.02, unpaired t test. All data are shown as mean ± SEM. \**p*<0.05, \*\**p*<0.01, \*\*\**p*<0.001.

With the decrease of histone acetylation by Smarcc2 deficiency, we next examined the potential interaction between Smarcc2 and histone deacetylases (HDACs) using co-immunoprecipitation (Co-IP) assays. As shown in **Fig. 6c**, Smarcc2 had a strong binding with HDAC2, and this association was decreased in Smarcc2^KD^ mice, which may free HDAC2 to deacetylate H3, resulting in reduced H3K9ac and H3K27ac.

The global reduction of histone acetylation by Smarcc2 deficiency prompted us to test whether synaptic genes had reduced histone acetylation at their promoters in Smarcc2^KD^ mice, which led to their transcriptional downregulation. To test this, we conducted chromatin immunoprecipitation (ChIP) assays to examine H3K9ac occupancy at the promotor region of *Slc1a3* (EAAT1), *Slc6a1* (GAT1), and *Slc32a1* (VGAT), representative synaptic genes with diminished expression in Smarcc2^KD^ mice. Primers were designed at the TSS region of these genes with strong H3K9ac occupancy from ChIP-seq landscapes (**Fig. 6d**). ChIP-PCR revealed a significant decrease of H3K9ac enrichment at *Slc1a3*, *Slc6a1*, and *Slc32a1* promoters in Smarcc2^KD^ mice, compared to controls (**Fig. 6e**), suggesting that Smarcc2 deficiency-induced downregulation of these synaptic genes may be attributable to the reduced histone acetylation.

### HDAC inhibitor restores some cognitive behaviors and synaptic genes in *Smarcc2*-deficient mice

With the identified reduction of histone acetylation by Smarcc2 knockdown, we next investigated whether elevating histone acetylation with HDAC inhibition could ameliorate phenotypes of Smarcc2^KD^ mice. The potent and brain-permeable class I HDAC inhibitor, romidepsin, was administered as we previously described (0.25 mg/kg, i.p. once daily for 3 days) [6, 24]. Immunostaining of H3K9ac demonstrated that the reduced H3K9ac fluorescence intensity in PFC of Smarcc2^KD^ mice was significantly restored following romidepsin treatment (**Fig. 7a**). Behavioral assays indicated that romidepsin treatment significantly improved the performance of Smarcc2^KD^ mice in spontaneous T maze working memory tests, as evidenced by the increased correctness across different trials (**Fig. 7b**). This rescuing effect was more pronounced in male Smarcc2^KD^ mice (**Fig. S4**). However, the spatial memory deficit in female Smarcc2^KD^ mice was not significantly improved by romidepsin treatment in BM tests (**Fig. 7c**).

**Figure 7.**
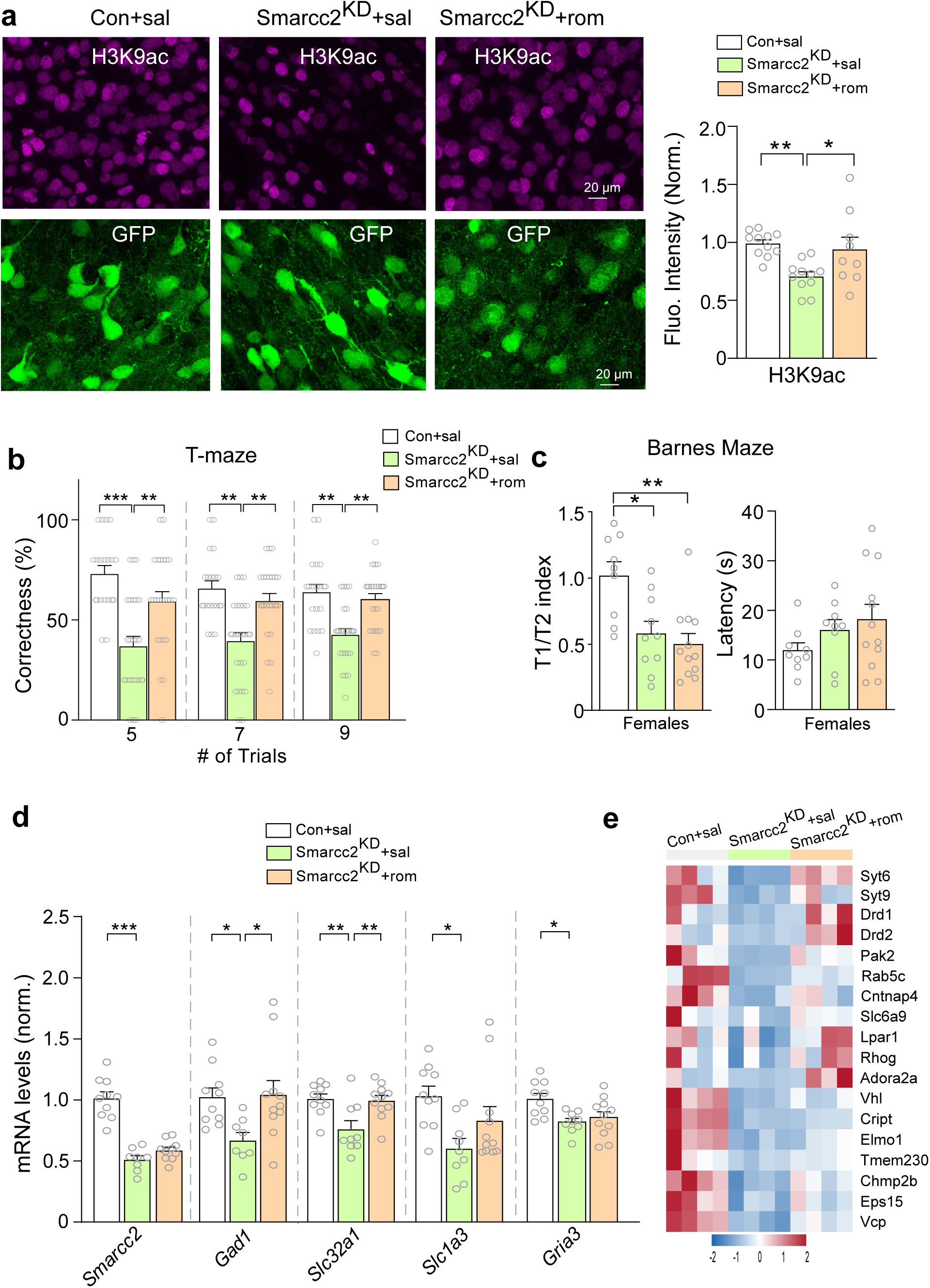
HDAC inhibitor treatment of Smarcc2^KD^ mice normalizes H3K9ac levels, rescues working memory deficits, and restores synaptic gene expression. **(a)** Representative confocal images (63×) showing H3K9ac immunostaining (magenta) and quantification of H3K9ac fluorescence intensity in control vs. Smarcc2^KD^ mice treated with saline or romidepsin. Scale bars: 20 μm. n=9-11 images/4 mice/group, F_2,28_=6.4, *p*=0.0051, one-way ANOVA. **(b)** Bar graph depicting the percentage of correct response in spontaneous T maze alternation tests of the 3 groups of mice. n=20-25 mice/group. Percent Correctness, 5 trials: F_2,56_=12.1, *p*<0.001, one-way ANOVA; 7 trials: KW=13.7, *p*=0.001; 9 trials: KW=15.1, *p*<0.001, K-W test. **(c)** Bar graphs showing the spatial memory index and latency to find the correct hole in BM tests of the 3 groups of female mice. n=9-12 mice/group. Spatial memory index: KW= 10.6, *p*=0.005, K-W test. **(d)** Bar graph showing the mRNA level of Smarcc2 and synaptic genes in the 3 groups of mice. n=8-11 mice/group. *Smarcc2*: F_2,25_=46.6, *p*<0.001; *Gad1*: F_2,26_=4.5, *p*=0.02; *Slc32a1*: F_2,27_=7.1, *p*=0.003; *Gria3*: F_2,25_=5.4, *p*=0.01, one-way ANOVA; *Slc1a3*: KW= 9.0, *p*=0.01, K-W test. **(e)** Heatmap showing the expression levels of synaptic genes in the 3 groups of mice from RNA-seq data. All data are shown as mean ± SEM, \**p*<0.05, \*\**p*<0.01, \*\*\**p*<0.001.

To find out potential mechanisms underlying the behavioral effects of romidepsin, we examined its impact on synaptic gene expression. As shown in qPCR profiling (**Fig. 7d**), while romidepsin treatment did not change *Smacc2*, it significantly or partially rescued several GABA- and glutamate-related genes diminished in Smarcc2^KD^ mice (*Gad1*, *Slc32a1,* and *Slc1a3*).

RNA-seq analysis further revealed that, among 653 downregulated genes by *Smarcc2* knockdown, 136 genes were elevated by romidepsin treatment (**Sup. Table 5**, 4.54 fold over-enrichment, hypergeometric p-value: 2.56E-53). Among these romidepsin-restored genes, 18 were synaptic genes (**Fig. 7e**), including *Syt6* and *Syt9* (encoding synaptotagmin 6 and 9) involved in Ca^2+^-dependent exocytosis of synaptic vesicles.

## Discussion

Smarcc2 is recognized as a key scaffold protein involved in the assembly and regulation of BAF chromatin remodeling complexes [25], which modulate genomic architecture by sliding or ejecting nucleosomes [26]. Disruption of BAF complexes has been frequently associated with human cancers [27] and developmental disorders [28]. Here, we found that Smarcc2 deficiency in PFC of adolescent mice led to the impaired working memory (T-maze) across both sexes and attenuated spatial memory (Barnes Maze) specifically in females. Working memory and spatial memory are fundamental cognitive functions [29]. These findings align with phenotypic observations in humans carrying *de novo* SMARCC2 mutations [16, 17], supporting the causal link of Smarcc2 haploinsufficiency to ASD/ID [2].

Accumulating evidence suggests that synaptic dysfunction is a key pathogenic factor in ASD [6, 9, 21, 30–32]. Our transcriptomic analyses revealed that Smarcc2 deficiency in PFC of adolescent mice significantly downregulated synaptic genes, particularly those involved in chemical synaptic transmission. Consistently, we found that embryonic (E16.5) cells from Smarcc1/2 double knockout mice [18] also had synaptic genes enriched in the downregulated gene set. In agreement with this, deep single-nucleus RNA sequencing of postmortem human brains revealed that the loss of synaptic genes in pre- and postsynaptic compartments of excitatory and inhibitory neurons was the most prominent change in ASD patients [33]. Quantitative PCR confirmed the reduction of inhibitory (*Gad1*, *Slc6a1*, *Slc32a1*) and excitatory (*Slc1a3*, *Gria3*) synaptic genes in Smarcc2-deficient mice. Electrophysiological recordings further revealed the significantly diminished GABAergic and glutamatergic synaptic currents in PFC pyramidal neurons of Smarcc2-deficient mice. These data have for the first time uncovered the critical role of Smarcc2 in maintaining synaptic function of mature neurons.

While BAF complexes are known to modulate chromatin structure, the precise epigenetic mechanisms by which Smarcc2 regulates synaptic genes remain unclear. We found that Smarcc2 bound to HDAC2, and Smarcc2 knockdown led to the decreased histone acetylation (H3K9ac and H3K27ac) without significantly affecting other histone marks (H3K27me3 and H3K4me3). H3K9ac and H3K27ac are associated with active transcription and play a pivotal role in development [34, 35]. Embryonic cells from Smarcc1/2 double knockout mice also had the reduced H3K9ac, however, a dramatic increase of H3K27me3, a gene repression marker [36], was thought to underlie the impact of Smarcc1/2 loss on genes involved in early development in these embryonic cells [18]. Our ChIP-PCR results on the reduced H3K9ac occupancy at promoters of downregulated synaptic genes in Smarcc2-deficient mice provide further evidence on the role of histone acetylation in Smarcc2 regulation of synaptic gene expression.

Smarcc2 knockdown reduced the interaction between Smarcc2 and HDAC2, which may lead to the enhanced association between HDAC2 and histone H3, causing the reduced histone acetylation. One therapeutic strategy for Smarcc2-deficient mice is to elevate histone acetylation via HDAC inhibition. Dysregulated HDAC activity has been implicated in both ASD and ID due to its impact on neurodevelopmental gene expression [37, 38]. HDAC inhibitors have been used in treating various ASD/ID models [5, 6, 39–41]. Here we found that a short treatment of Smacc2-deficient mice with romidepsin, a selective class I HDAC inhibitor, restored H3K9ac levels, improved working memory, and normalized synaptic genes expression and synaptic function. These findings underscore the therapeutic potential of HDAC inhibitors in ameliorating phenotypes associated with Smarcc2 haploinsufficiency.

Collectively, based on our data, we propose a model revealing the link of Smarcc2 to ASD/ID (**Fig. 8**). Smarcc2 deficiency in PFC leads to the reduced expression of GABA- and glutamate-related synaptic genes via decreased histone acetylation, resulting in the impaired inhibitory and excitatory synaptic transmission, and cognitive performance. These synaptic and behavioral deficits can be ameliorated by HDAC inhibition, providing a therapeutic strategy for Smarcc2-associated neurodevelopmental disorders.

**Figure 8.**
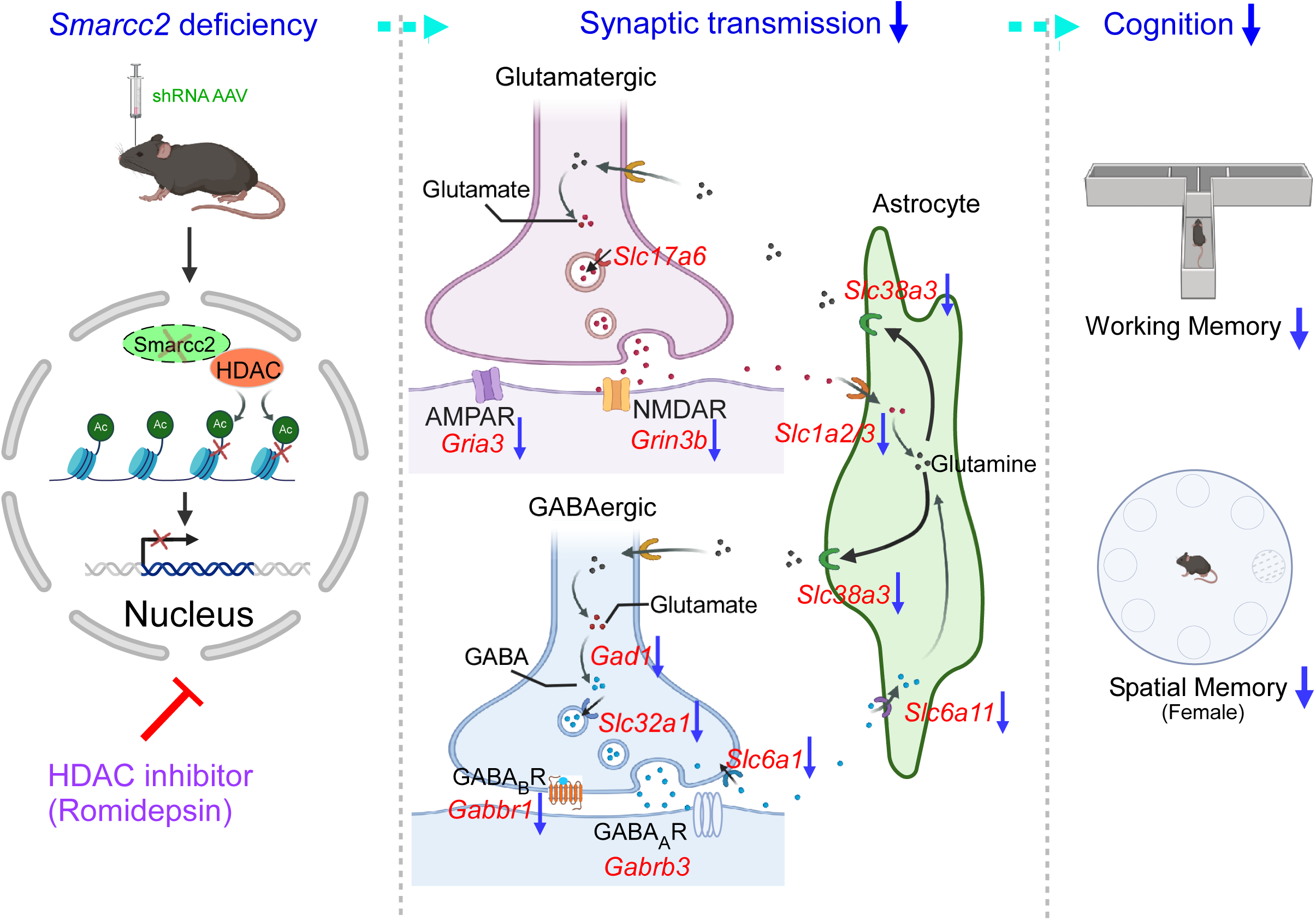
A schematic model illustrating the impact of Smarcc2 deficiency and a therapeutic strategy. Knockdown of Smarcc2, a core subunit of chromatin remodeler BAF complex, decreases the level of histone acetylation, leading to the reduced expression of synaptic transmission-related genes in both glutamatergic and GABAergic synapses. The resulting attenuation of excitatory and inhibitory synaptic transmission leads to cognitive deficits (WM, BM). Treatment with the class I HDAC inhibitor romidepsin restores H3K9ac levels and rescues synaptic genes expression and synaptic function, leading to cognitive improvement.

## Materials and Methods

### Generation and analysis of human iPSC-derived neurons

Human induced pluripotent stem cells (iPSCs) from both neurotypical controls and idiopathic ASD patients (4 lines per group with each group comprising 2 males and 2 females, 4-10 years old at sampling) were acquired from the Human Pluripotent Stem Cell Line Repository at the California Institute for Regenerative Medicine (CIRM). These iPSCs were generated using non-integrating Episomal vectors. All iPSCs were maintained on gamma-irradiated CF-1 mouse embryonic fibroblasts in DMEM/F12 medium supplemented with knockout serum replacement, β-mercaptoethanol, NEAA, L-glutamine, and FGF2. The human iPSCs were differentiated to cortical neurons as we described [42]. Briefly, iPSCs were dissociated to form embryoid bodies (EBs), cultured in suspension in a DMEM/F12 and Neurobasal mix with supplements including N2, B27 (vitamin A-free), NEAA, ascorbic acid, SB431542, dorsomorphin, XAV939, and Cyclopamine. On day 6, EBs were plated on Matrigel-coated plates and maintained for 6 days. At day 12, dorsal cortical progenitors were dissociated with Accutase and plated on polyornithine/Matrigel-coated plates at 5,000–10,000 cells/cm² in the same medium with Y27632 added for the first 24 hours. Cells were passaged again on day 16. From day 18, the medium was changed to Neurobasal with B27 (vitamin A-free), BDNF, GDNF, dcAMP, ascorbic acid, and DAPT to promote cortical neuron differentiation. Half-medium changes were performed every other day, and neurons were cultured until day 60 for experimental use. Immunostaining and qPCR were performed using protocols consistent with those for mouse tissues.

### Animals and Reagents

All experiments were conducted in accordance with the guidelines of the institutional animal care and use committee (IACUC) in State University of New York at Buffalo. Five-week-old C57BL/6J mice (Strain #: 000664, The Jackson Lab) were used in this study. To knockdown *Smarcc2*, a short hairpin RNA (shRNA) sequence (CCCGATAGTTGATCCTGAGAA) was cloned into GFP-tagged adeno-associated virus (AAV) vector (Addgene) under the control of U6 promotor. Viral particles (8×10^14^ vg/ml) were produced by the viral core center of Emory University. Romidepsin (rom, Selleckchem, S3020) was prepared in DMSO and stored at −20 °C at the stock concentration of 1mg/ml. Prior to use, the stock solution was diluted with saline.

### Animal Surgery

In brief, mice were anaesthetized with ketamine/xylazine. Stereotaxic AAV injections were performed in the medial PFC (2.0 mm anterior to bregma; 0.3 mm lateral; 2.0 mm dorsal to ventral; 1 μl per hemisphere) [9]. The injections were administered with a Hamilton syringe (gauge 34) at a speed of 0.1 μl/min.

### Behaviors

All behavioral tests were conducted 2 weeks post-stereotaxic injection. All the tests were performed between 10 a.m. and 6 p.m. under dim lighting conditions. Mice were acclimated to the testing room for 1h before tests. Any-maze software was used to track and record behavioral data. To minimize olfactory interference, testing equipment was cleaned with 70% ethanol between different animals. Additionally, to avoid the potential disturbance of gender, male and female mice were tested separately.

*<u>Spontaneous T maze</u>* was used to assess working memory, with modification based on the protocol [43]. The T-shaped maze consists of a start area (12’’Lx5.5’’Wx6’’H) and two arms (23’’Lx5.5’’Wx6”H). Mice were initially placed in the start area for 30s, after which they were allowed to choose either arm within 2 min. Upon complete entry into an arm, the mouse was confined within that arm for 30s before being returned to the start area. If a mouse failed to exit an arm within 10s, a clean paper was used to gently guide it out. The procedure was repeated for 10 trials. A correct choice was defined as selecting an arm opposite to the chosen one in the previous trial. The percentage of correctness was calculated from trial 2 to trail 10.

*<u>Barnes Maze (BM)</u>* was employed to evaluate the spatial memory. As described previously [44], mice were placed at the edge of a round arena containing 8 evenly spaced holes. Three different visual cues were placed on the surrounding walls. The mouse faced the opposite direction of the correct hole, which has an escape box attached underneath. A bright light overhead was used as an aversive stimulus. After 5-min habituation, the mouse underwent two training sessions, each lasting 5 min, with a 5-min rest interval in the holding cage between sessions. After 15 min break, the mouse was put back on the arena for the final test in which the escape box was removed. Time spent exploring the correct hole (T1) and incorrect holes (T2) were recorded. T1/T2 index was calculated to evaluate the spatial memory.

*<u>Social preference test</u>* was performed to assess sociability. As described previously [45], a three-chamber apparatus with two inverted cups in the side chambers was used. One day before the test, the mouse was put in the center chamber and allowed to explore the three chambers with two empty cups for 10 min (habituation). On the test day, two identical non-social objects (paper balls) were placed underneath the inverted cups in the three-chamber apparatus for the mouse to explore (10 min). Following a 5-min rest in the holding cage, the mouse was put back in the three chambers with two cups containing one non-social object (wooden block) and one social object (sex- and age-matched mouse) to explore for another 10 min. The time interacting with social (T_soc_) and non-social objects (T_NS_) were recorded. (T_soc_-T_NS_)/ (T_soc_+T_NS_) was calculated as social preference index.

*<u>Elevated plus maze (EPM)</u>* was utilized to assess anxiety. As described preciously [44], mice were placed at the center of a plus-shaped arena, facing an open arm and allowed to explore freely for 10 min. Time spent in the open arms and closed arms, latency to the open arms, and the total distance traveled were recorded.

*<u>Open Field (OF)</u>* was conducted to assess anxiety and locomotor activity. As described previously [44], the mouse was placed in the center of a rectangular arena with opaque walls and allowed to explore freely for 10 min. The center area was defined as half length and width of the arena. Total distance traveled and time spent in the center area were recorded.

*<u>Light and dark box test</u>* was used to evaluate anxiety [46]. The apparatus consisted of a light and a dark chamber of equal size (14.5”Lx10.5”Wx13”H). Mice were placed in the light box, facing away from the door. The time spent in the light box and the frequency to enter the light box within 10 min was recorded.

*<u>Grooming</u>* was used to test repetitive behavior [47]. The mouse was put in a holding cage individually and allowed to explore for 30 min. A camera was used to record the whole session. The grooming time and frequency were counted manually for the last 10 min.

*<u>Resident-intruder (RI) test</u>* was used to evaluate aggression [24], Briefly, mice were single housed for 1 day before test. During the test, an intruder (an unfamiliar, group-housed, same sex WT mouse of 5-15% lighter) was introduced into the resident’s home cage for 10 min. Each intruder was used only once. Aggressive behavior of the resident mouse against the intruder was scored. The duration and frequency of aggression, as well as the total social interactions between the resident mouse and the intruder, were counted manually.

### Western blot

Nuclear protein extraction was performed as we previously described [9] with modifications. Briefly, GFP-positive PFC tissue was dissected under a microscope with fluorescence adapter system (Avantor Science Central, 77463-320). Tissues were homogenized with 250 μl hypotonic buffer with protease inhibitor cocktail and 1 mM PMSF, followed by incubation on ice for 15 min. Subsequently, 12.5 μl of 10% NP-40 was added and the mixture was vortexed for 10 s. The homogenate was centrifuged at 3000 g for 10 min at 4 °C. The nuclear pellet was resuspended in filtered 1% SDS containing protease inhibitor cocktail and 1 mM PMSF. All samples were normalized to the same concentration and boiled in 4x SDS loading buffer for 10 min. Proteins were separated using 6% and 12% SDS-polyacrylamide gels. A total of 10 μg proteins was loaded for target proteins, while 5 μg was used for Histone H3. Western blotting was performed using antibodies against Smarcc2 (Cell Signaling, 12760, 1:1000), H3K9ac (Cell Signaling, 9649, 1:1000), H3K27ac (Cell Signaling, 8173, 1:1000), H3K27me3 (Cell Signaling, 9733, 1:1000) and Histone H3 (Cell Signaling, 4499, 1:1000).

### Real-time RT-PCR

Total RNA was extracted from GFP positive mouse PFC using Trizol reagent (Invitrogen, 15596026). cDNA was synthesized using the iScriptTM cDNA synthesis Kit (Bio-Rad, 1708891). Quantitative real-time PCR was performed using the CFX Connect Real-Time PCR Detection System and iQ™ SYBR® Green Supermix (Bio-Rad, 1708882). GAPDH was used as the housekeeping gene for normalization. Each 20 μl reaction was run in a 96-well PCR plate (Bio-Rad, HSP9601) using the following cycling parameters: 95 °C for 3 min followed by 40 cycles of 95 °C for 15 s, 60 °C for 20 s, and 72 °C for 30 s. Fold changes in the target gene expression were calculated as following: ΔCt =Ct(target) – Ct(GAPDH), and Δ(ΔCt) =ΔCt(Smarcc2^KD^) – mean ΔCt(control) or Δ(ΔCt) =ΔCt (Smarcc2^KD^+drug) – mean ΔCt(Smarcc2^KD^+saline), and Fold change = 2^-Δ(ΔCt)^. Primers used in qPCR experiments are included in **Sup. Table 6**.

### Immunohistochemistry

Mice were anesthetized with ketamine/ xylazine and transcardially perfused with phosphate buffered saline (PBS) and 4% fresh prepared paraformaldehyde (PFA) [9]. Brains were post-fixed in 4% PFA overnight, dehydrated in 30% sucrose for 2 days and sectioned coronally (50 μm thickness). After three washes with PBS (10 min each), slices were blocked with 5 % BSA containing 0.3% Triton for 2 hr at room temperature. Then slices were incubated overnight at 4 °C with primary antibody against Smarcc2 (Cell Signaling, 12760, 1:1000), H3K9ac (Cell Signaling, 9649; 1:1000) or NeuN (Millipore, MAB377; 1:250). Following three PBS washes (15 min each), slices were incubated with secondary antibodies: Alexa Fluor donkey anti mouse-568 (Invitrogen, A10037; 1:1000) and Alexa Fluor goat anti rabbit-647 (Invitrogen, A21245; 1:1000) for 2 hr at room temperature. Nuclei were counterstained with DAPI (ThermoFisher, D1306, 5 μg/ml) for 30 min at room temperature. Sections were mounted using ProLong™ Diamond Antifade Mountant (Invitrogen, P36970). Images were acquired using 63× objective with a Leica TCS SP8 confocal microscope. Identical imaging conditions and parameters were applied to all specimens using Image J software (version 1.52p, NIH).

### Electrophysiological recordings of brain slices

The whole-cell voltage-clamp recording technique was used to measure synaptic currents in slices as previously described [48]. Mice were decapitated after inhaling isoflurane and the brain was quickly removed, iced, and coronally cut into 300 µm slices with a vibratome (Leica VP1000S, Leica Microsystems Inc, Germany) in an ice-cold sucrose solution. The slices were incubated in artificial cerebrospinal fluid (ACSF; in mM: 130 NaCl, 26 NaHCO_3_, 1 CaCl_2_, 5 MgCl_2_, 3 KCl, 1.25 NaH_2_PO_4_, 10 glucose, pH 7.4, 300 mOsm) in 30-35 °C for 1 hr and maintained in room temperature (21 °C), bubbling with 95% O_2_ and 5% CO_2_. The mouse slices were transferred in a perfusion chamber attached to the fixed stage of an upright microscope (Olympus) and submerged in continuously flowing oxygenated ACSF. Layer V mPFC pyramidal neurons were visualized with a 40X water-immersion lens and recorded with the Multiclamp 700 A amplifier (Molecular Devices, Sunnyvale, CA), Clampex software 9 (Molecular Devices, Sunnyvale, CA) and Digidata 1322A (Molecular Device, Sunnyvale, CA, USA). Recording pipette contained the following internal solution (in mM: 130 Cs-methanesulphonate, 10 CsCl, 4 NaCl, 1 MgCl_2_, 10 HEPES, 5 EGTA, 2 QX-314, 12 phosphocreatine, 5 MgATP, 0.5 Na_2_GTP, pH 7.2–7.3, 265–270 mOsm).

For spontaneous inhibitory postsynaptic current (sIPSC) recording, neurons were held at 0 mV, for excitatory postsynaptic current (sEPSC) recording, neurons were held at −70 mV. Evoked GABA_A_R-IPSC (at 0 mV) and AMPAR-EPSC (at −70 mV) were generated with a series of pulses generated by S48 pulse stimulator (Grass Technologies, West Warwick, RI, USA) with different stimulation intensities (0.06 ms, 50–90 μA) delivered at 0.1 Hz. A bipolar stimulating electrode (FHC, Bowdoinham, ME) was placed ∼100 μm from the neuron under recording. For paired-pulse ratios, GABA_A_R-IPSC was evoked by double pulses with a 30ms, 50ms, or 100ms interval, AMPAR-EPSC was evoked with 20ms, 50ms, or 100ms interval.

### Co-immunoprecipitation (Co-IP)

As described in previous paper [7], transfected N2A cells were collected and homogenized with cold RIPA lysis buffer (mM: 150 NaCl, 50 Tris (pH=8.0), 1% Nonidet P-40, 0.5% sodium deoxycholate, and 0.1% SDS) containing protease inhibitor. The lysates were incubated on ice for 30 min and then centrifuged at 12,000 g for 15 min at 4 °C. Supernatant fraction (40 μl) was collected as input control. The remainder was pre-cleared with Protein A/G plus agarose (Santa Cruz Biotechnology, sc-2003, 100 μl) for 1 hr at 4 °C. The pre-cleared supernatant was incubated overnight at 4 °C with antibodies against HDAC2 (Abcam, ab12169, 10 μg) or mouse IgG (sigma, 12-371, 10 μg) as negative control. Following incubation with 60 μl protein A/G plus agarose for 2 hr at 4 °C, immunoprecipitates were washed three times with lysis buffer and then boilded in 2× SDS loading buffer (40 μl). Western blotting was performed using antibodies against Smarcc2 (Cell Signaling, 12760, 1:1000), HDAC1 (Cell Signaling, 34589, 1:1000), and HDAC2 (Abcam, ab12169, 1:1000).

### Chromatin immunoprecipitation (ChIP) assay

As previously described [9], GFP positive PFC from three mice were pooled per sample (50 mg in total). Each sample was homogenized in 250 μl ice cold douncing buffer with 1 ml 26-gauge syringe (15 strokes). Chromatin was digested using micrococcal nuclease (5 U/ml, Sigma, N5386) for 7 min at 37 °C, then the reaction was terminated by EDTA (10 mM, Invitrogen, 15575-038). Following lysis in 1 ml hypotonic lysis buffer (0.2 mM EDTA, pH 8.0, 0.1 mM benzamidine, 0.1 mM PMSF, 1.5 mM DTT) containing protease inhibitor on ice for 1 hr, lysates were centrifuged at 3000 g for 5 min at 4 °C. Supernatant (2-5 %) was reserved as input control, while the remainder was pre-cleared with Salmon sperm DNA/protein A agarose-50% slurry (Millipore, 16–157) for 2 hr at 4 °C. Pre-cleared chromatin was incubated with antibodies against H3K9ac (10 μg per reaction; Sigma, 17-658) or Rabbit IgG (10 μg per reaction, Sigma, PP64B) overnight at 4 °C, followed by protein A agarose incubation for 2 hr. After washing, immunoprecipitates were eluted in 1% SDS, 0.1 M NaHCO_3_. Crosslinks were reversed by incubation with 5M NaCl (8 μl), 0.5 M EDTA (4 μl), 1M Tris-HCl (pH 7.4, 8 μl), and 20 mg/ml proteinase K (0.4 μl, Thermo Fisher, EO0491) at 50 °C for 1 hr. DNA was purified using the QIAquick PCR purification Kit (Qiagen, 28104) and eluted in 30 μl Buffer EB (Qiagen, 19086). ChIP-PCR was performed using primers targeting the promoter region of *Slc1a3* (Forward, 5’-GAGGACTGGGCTTGCATAGTT-3’; Reverse, 5’-TTCCTCCCATCCCAGAGTCA-3’), *Slc6a1* (Forward, 5’-AGCCCTAGAGAGCTGAGAGGTT-3’; Reverse, 5’-GAGGACGGGGGACGATGC-3’), *Slc32a1* (Forward, 5’-CAAGAGCCAGACTGTCGTGA-3’; Reverse, 5’-ACTTGTTGGACACGGAGGTG-3’). Quantification of ChIP signals was calculated as % input.

### RNA sequencing and data analysis

Human iPSC-derived neurons (2 groups, 4 lines/group) and mouse brain tissues (3 groups, 4 samples/group, dissected GFP+ PFC pooled from 3 mice (50 mg)/sample) were used for RNA-seq. Total RNA was extracted using the same method as described in the real-time RT-PCR section. RNA-seq libraries were prepared using the TruSeq Stranded Total RNA Plus Ribo-Zero Kit (Illumina) and sequenced on the HiSeq 2500 platform (Illumina) at LC Sciences. Raw paired-end FASTQ reads were cleaned with cutadapt (v. 2024.10.0) and aligned to the *Mus musculus* reference genome (mm10) using HISAT2 (v. 2.2.1). Transcripts assembly of each individual sample, merging and gene level expression counts were quantified using StringTie (v. 2.2.3) and offcompare (v. 0.12.6). Differential gene expression analysis was conducted using DESeq2 (v. 3.2). For human iPSC-Neuron RNA-seq data, DEGs were identified based on *P* ≤ 0.05 relative to controls. For mouse RNA-seq data, DEGs were identified based on a fold change (FC) ≥ 1.2 and *P_adj._* ≤ 0.05 relative to controls.

Biological Processes Gene ontology (GO) analysis was performed using EnrichR (https://maayanlab.cloud/Enrichr/). Overlap of genesets was done using InteractiVenn (https://www.interactivenn.net/). Functional protein classification of top GO categories was conducted using String database (https://string-db.org/) and exported into CytoScape (v. 3.10.3) to generate the protein-protein interaction plot. Genes were classified into categories (when applicable) with the String-db’s Cytoscape (StringApp) plugin. Hub genes were identified, and interactome networks were generated using CytoHubba (v. 0.1) with the maximum clique centrality method. Volcano plots, heatmaps and GO bar plots were generated using ggplot2 (v. 3.5.1). Synaptic specific gene ontology was done using SynGO exported onto a sunburst plot (https://www.syngoportal.org/).

### ChIP-seq data analysis

Smarcc2 ChIP-seq data consisting of raw fastq files were obtained via Gene Expression Omnibus (GEO) under the accession number GSM5311593, GSM4837715, GSM4837716, GSM5311594, and GSM5311596. Reads were mapped to the mouse reference genome mm10 using Bowtie2 (v. 2.4.2) with default parameters. Mapped bam files were then filtered using samtools with a mapping quality (MAPQ) score cutoff of 20. Peak calling was then performed with MACS2 (v. 2.1.1.2+) with a p≤0.05 cutoff for peak detection otherwise under default settings. Peaks were then annotated with ChIPseeker (v. 1.8.0). To visualize genomic coverage, bigWig files were generated with bamCompare (v. 3.3.2.0.0) by deepTools with bin size set to 50bp. Bigwig files were then visualized in the Interactive Genome Browser (IGV: https://igv.org/) for further examination. These data were then analyzed with additional deepTools packages, in which bigWig files were prepared in computeMatrix (v. 3.1.2.0.0) for genomic regions within 1kb of the TSS and plotted using plotHeatmap (v3.1.2.0.1). Pathway analysis was performed using EnrichR on peaks within 1 kb of TSS.

### Statistical analysis

Statistical analysis was performed with GraphPad Prism 8.4.3. Data normality was assessed using the Shapiro-Wilk test before parametric analyses. Comparisons between two independent groups were conducted using two-tailed unpaired t-tests for normally distributed data, otherwise, the Mann-Whitney (M-W) test was applied. For comparisons among more than two groups, one-way ANOVA was used for normally distributed data with homogeneity of variance, followed by Bonferroni *post hoc* tests for multiple comparisons, otherwise Kruskal-Wallis (K-W) test was performed, followed by pairwise Wilcoxon rank-sum tests with Bonferroni correction. In the social preference test, social versus nonsocial time was analyzed using two-way ANOVA. All data are presented as mean ± SEM. All experiments were replicated in multiple cohorts of animals and at least three independent *in vitro* experiments.

## Supporting information

All Supplementary Figures

All Supplementary Tables

## Acknowledgements

This work was supported by a grant from the National Institutes of Health NS127728 to Z.Y. and J.F.

## Author Contributions

P.L. performed animal surgery, immunostaining, behavioral, qPCR, and biochemical experiments, and wrote the draft. S.M. performed animal surgery and electrophysiological experiments. P.J. P. and Z.Y. performed some bioinformatics analysis. P.Z. performed earlier electrophysiological experiments. K.S., K.W.T. and J.F. performed experiments with human iPSC-derived neurons. Z.Y. supervised the project and edited the paper.

## References

1. Shenouda, J., et al., Prevalence and Disparities in the Detection of Autism Without Intellectual Disability. Pediatrics, 2023. 151(2).

2. Satterstrom, F.K., et al., Large-Scale Exome Sequencing Study Implicates Both Developmental and Functional Changes in the Neurobiology of Autism. Cell, 2020. 180(3): p. 568–584.e23.

3. Sanders, S.J., et al., De novo mutations revealed by whole-exome sequencing are strongly associated with autism. Nature, 2012. 485(7397): p. 237–41.

4. Vissers, L.E., C. Gilissen, and J.A. Veltman, Genetic studies in intellectual disability and related disorders. Nat Rev Genet, 2016. 17(1): p. 9–18.

5. Yan, Z., Targeting epigenetic enzymes for autism treatment. Trends Pharmacol Sci, 2024. 45(9): p. 764–767.

6. Qin, L., et al., Social deficits in Shank3-deficient mouse models of autism are rescued by histone deacetylase (HDAC) inhibition. Nat Neurosci, 2018. 21(4): p. 564–575.

7. Wang, Z.J., et al., Amelioration of autism-like social deficits by targeting histone methyltransferases EHMT1/2 in Shank3-deficient mice. Mol Psychiatry, 2020. 25(10): p. 2517–2533.

8. Katayama, Y., et al., CHD8 haploinsufficiency results in autistic-like phenotypes in mice. Nature, 2016. 537(7622): p. 675–679.

9. Qin, L., et al., Deficiency of autism risk factor ASH1L in prefrontal cortex induces epigenetic aberrations and seizures. Nat Commun, 2021. 12(1): p. 6589.

10. Herrera, M.L., et al., Targeting epigenetic dysregulation in autism spectrum disorders. Trends Mol Med, 2024. 30(11): p. 1028–1046.

11. Lessard, J., et al., An essential switch in subunit composition of a chromatin remodeling complex during neural development. Neuron, 2007. 55(2): p. 201–15.

12. He, S., et al., Structure of nucleosome-bound human BAF complex. Science, 2020. 367(6480): p. 875–881.

13. Fu, J.M., et al., Rare coding variation provides insight into the genetic architecture and phenotypic context of autism. Nature Genetics, 2022. 54(9): p. 1320–1331.

14. Barisic, D., et al., Mammalian ISWI and SWI/SNF selectively mediate binding of distinct transcription factors. Nature, 2019. 569(7754): p. 136–140.

15. Clapier, C.R., et al., Mechanisms of action and regulation of ATP-dependent chromatin-remodelling complexes. Nat Rev Mol Cell Biol, 2017. 18(7): p. 407–422.

16. Machol, K., et al., Expanding the Spectrum of BAF-Related Disorders: De Novo Variants in SMARCC2 Cause a Syndrome with Intellectual Disability and Developmental Delay. Am J Hum Genet, 2019. 104(1): p. 164–178.

17. Bosch, E., et al., Elucidating the clinical and molecular spectrum of SMARCC2-associated NDD in a cohort of 65 affected individuals. Genet Med, 2023. 25(11): p. 100950.

18. Narayanan, R., et al., Loss of BAF (mSWI/SNF) Complexes Causes Global Transcriptional and Chromatin State Changes in Forebrain Development. Cell Rep, 2015. 13(9): p. 1842–54.

19. Tuoc, T., et al., Ablation of BAF170 in Developing and Postnatal Dentate Gyrus Affects Neural Stem Cell Proliferation, Differentiation, and Learning. Mol Neurobiol, 2017. 54(6): p. 4618–4635.

20. Reinert, S., et al., Mouse prefrontal cortex represents learned rules for categorization. Nature, 2021. 593(7859): p. 411-417.

21. Yan, Z. and B. Rein, Mechanisms of synaptic transmission dysregulation in the prefrontal cortex: pathophysiological implications. Mol Psychiatry, 2022. 27(1): p. 445–465.

22. Stoner, R., et al., Patches of disorganization in the neocortex of children with autism. N Engl J Med, 2014. 370(13): p. 1209–1219.

23. Hyun, K., et al., The BAF complex enhances transcription through interaction with H3K56ac in the histone globular domain. Nature Communications, 2024. 15(1): p. 9614.

24. Hernandez Carballo, L.G., et al., Systemic histone deacetylase inhibition ameliorates the aberrant responses to acute stress in socially isolated male mice. J Physiol, 2024. 602(9): p. 2047–2060.

25. Mashtalir, N., et al., Modular Organization and Assembly of SWI/SNF Family Chromatin Remodeling Complexes. Cell, 2018. 175(5): p. 1272–1288.e20.

26. Wilson, B.G. and C.W. Roberts, SWI/SNF nucleosome remodellers and cancer. Nat Rev Cancer, 2011. 11(7): p. 481–92.

27. Kadoch, C. and G.R. Crabtree, Mammalian SWI/SNF chromatin remodeling complexes and cancer: Mechanistic insights gained from human genomics. Sci Adv, 2015. 1(5): p. e1500447.

28. Valencia, A.M., et al., Landscape of mSWI/SNF chromatin remodeling complex perturbations in neurodevelopmental disorders. Nat Genet, 2023. 55(8): p. 1400–1412.

29. Murray, J.D., J. Jaramillo, and X.J. Wang, Working Memory and Decision-Making in a Frontoparietal Circuit Model. J Neurosci, 2017. 37(50): p. 12167–12186.

30. Bourgeron, T., From the genetic architecture to synaptic plasticity in autism spectrum disorder. Nat Rev Neurosci, 2015. 16(9): p. 551–63.

31. Cho, H., et al., *Adnp-mutant mice with cognitive inflexibility, CaMKII*α *hyperactivity, and synaptic plasticity deficits*. Mol Psychiatry, 2023. 28(8): p. 3548–3562.

32. Jung, E.M., et al., Arid1b haploinsufficiency disrupts cortical interneuron development and mouse behavior. Nat Neurosci, 2017. 20(12): p. 1694–1707.

33. Wamsley, B., et al., Molecular cascades and cell type-specific signatures in ASD revealed by single-cell genomics. Science, 2024. 384(6698): p. eadh2602.

34. Qiao, Y., et al., Dual roles of histone H3 lysine 9 acetylation in human embryonic stem cell pluripotency and neural differentiation. J Biol Chem, 2015. 290(4): p. 2508–20.

35. Creyghton, M.P., et al., Histone H3K27ac separates active from poised enhancers and predicts developmental state. Proc Natl Acad Sci U S A, 2010. 107(50): p. 21931–6.

36. Cai, Y., et al., H3K27me3-rich genomic regions can function as silencers to repress gene expression via chromatin interactions. Nature Communications, 2021. 12(1): p. 719.

37. Tseng, C.J., et al., Epigenetics of Autism Spectrum Disorder: Histone Deacetylases. Biol Psychiatry, 2022. 91(11): p. 922–933.

38. Ramaswami, G., et al., Integrative genomics identifies a convergent molecular subtype that links epigenomic with transcriptomic differences in autism. Nat Commun, 2020. 11(1): p. 4873.

39. Rein, B., et al., Inhibition of histone deacetylase 5 ameliorates abnormalities in 16p11.2 duplication mouse model. Neuropharmacology, 2022. 204: p. 108893.

40. Sun, W., et al., Histone Acetylome-wide Association Study of Autism Spectrum Disorder. Cell, 2016. 167(5): p. 1385–1397.e11.

41. Wang, W., et al., Histone Deacetylase Inhibition Restores Behavioral and Synaptic Function in a Mouse Model of 16p11.2 Deletion. Int J Neuropsychopharmacol, 2022. 25(10): p. 877–889.

42. Jiang, H., et al., Generation of human induced pluripotent stem cell-derived cortical neurons expressing the six tau isoforms. J Alzheimers Dis, 2025: p. 13872877251334831.

43. Deacon, R.M.J. and J.N.P. Rawlins, T-maze alternation in the rodent. Nature Protocols, 2006. 1(1): p. 7–12.

44. Li, P. and Z. Yan, An epigenetic mechanism of social isolation stress in adolescent female mice. Neurobiol Stress, 2024. 29: p. 100601.

45. Rein, B., K. Ma, and Z. Yan, A standardized social preference protocol for measuring social deficits in mouse models of autism. Nat Protoc, 2020. 15(10): p. 3464–3477.

46. Kulesskaya, N. and V. Voikar, Assessment of mouse anxiety-like behavior in the light-dark box and open-field arena: role of equipment and procedure. Physiol Behav, 2014. 133: p. 30–8.

47. Conrow-Graham, M., et al., A convergent mechanism of high risk factors ADNP and POGZ in neurodevelopmental disorders. Brain, 2022. 145(9): p. 3250–3263.

48. Tan, T., et al., Stress Exposure in Dopamine D4 Receptor Knockout Mice Induces Schizophrenia-Like Behaviors via Disruption of GABAergic Transmission. Schizophr Bull, 2019. 45(5): p. 1012–1023.

