## Supplementary material for "Cognitive and Synaptic Impairment Induced by Deficiency of Autism Risk Gene *Smarcc2* and its Rescue by Histone Deacetylase Inhibition": All Supplementary Figures

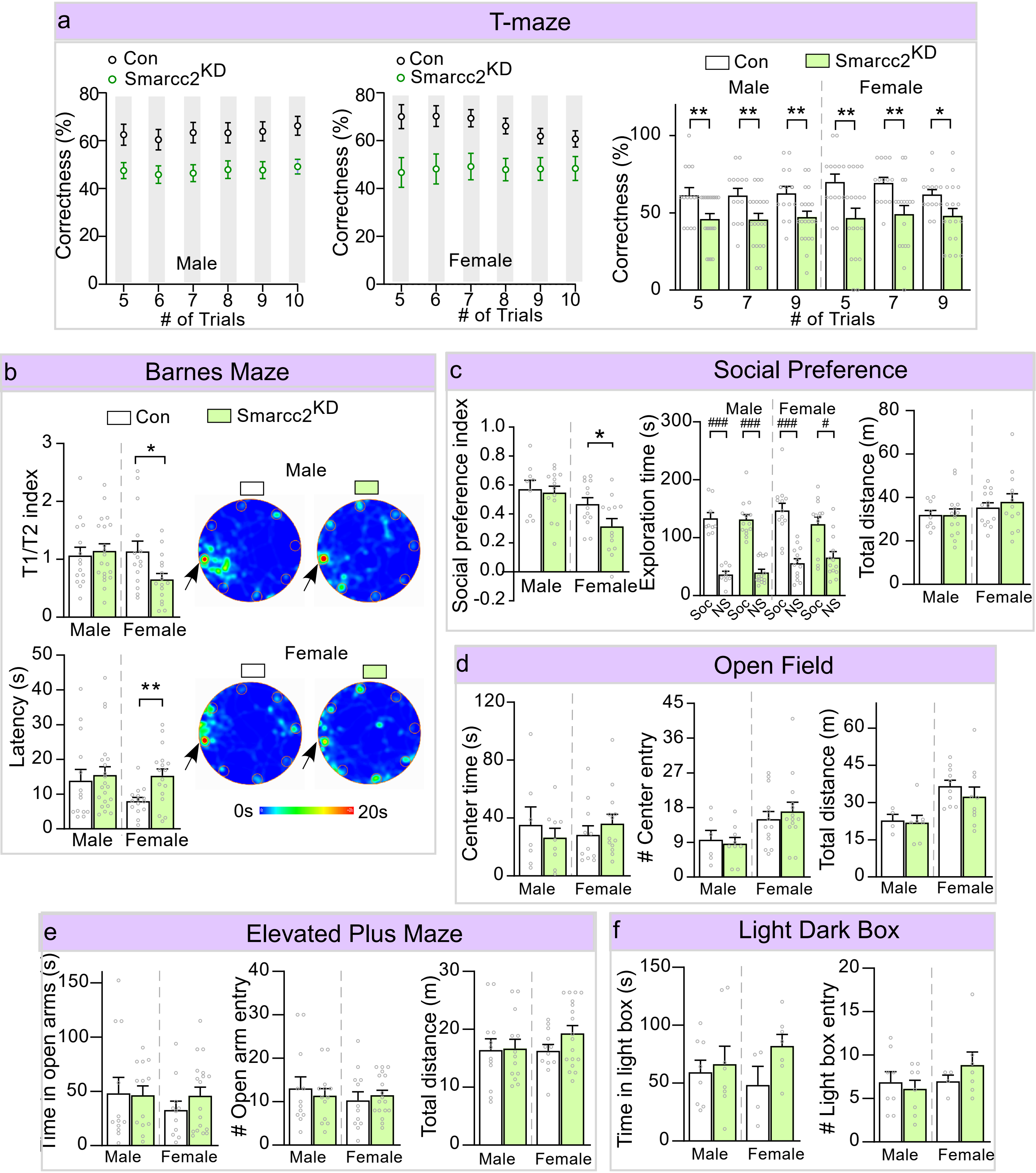


**Figure S1. Smarcc2 knockdown impairs working memory in both sexes and spatial memory in females. (a)** Plots showing the percentage of correct responses across trials in T-maze tests of control vs. Smarcc2$^{KD}$ mice of each sex. Con: n=16M, 14F, Smarcc2$^{KD}$: n=23M, 18F. **(b)** Bar graphs showing the spatial memory index and latency to find the correct hole in Barnes Maze (BM) tests of control vs. Smarcc2$^{KD}$ mice of each sex. Con: n=15M, 15F, Smarcc2$^{KD}$: n=20M, 16F. **(c)** Bar graphs showing the social preference index, time on social (Soc) and non-social (NS) objects, and total distance in three-chamber social preference tests of control vs. Smarcc2$^{KD}$ mice of each sex. Con: n=9M, 13F, Smarcc2$^{KD}$: n=14M, 13F. **(d)** Bar graphs showing the time spent in center area of open field (OF), the number of entries to center, and the total distance traveled in OF tests of control vs. Smarcc2$^{KD}$ mice of each sex. Con: n=7M, 11F, Smarcc2$^{KD}$: n=9M, 13F. **(e)** Bar graphs showing the time in open arm, number of entries to open arms and the total distance in EPM tests of control vs. Smarcc2$^{KD}$ mice of each sex. Con: n= 12M, 12F, Smarcc2$^{KD}$: n=13M, 18F. **(f)** Bar graphs showing the time spent in light box and the entry numbers to the light box in light-dark box tests of control vs. Smarcc2$^{KD}$ mice of each sex. Con: n=8M, 4F, Smarcc2$^{KD}$: n=8M, 7F. All data are shown as mean ± SEM, *$p<0.05$, **$p<0.01$, ***$p<0.001$. Detailed statistical data are provided in Source Data files.

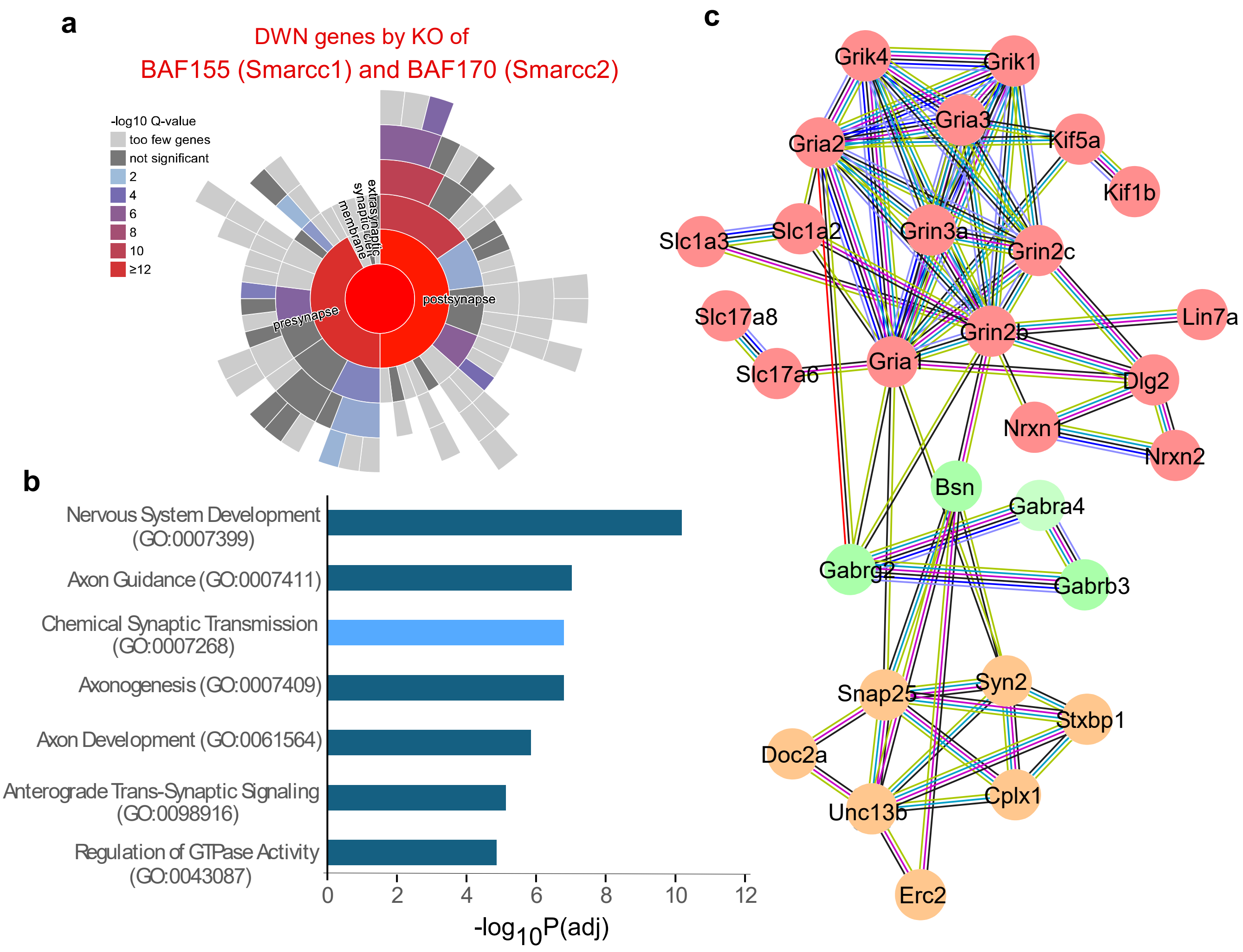


**Figure S2. Synaptic genes are downregulated in embryonic cells with double knockout of Smarcc1 and Smarcc2.(a)** Sunburst plot from SynGO analysis illustrating the most enriched cellular component categories for downregulated genes in E16.5 cells from Smarcc1/Smarcc2 double knockout mice. Higher red intensities indicate stronger enrichment. **(b)** GO analysis showing the most enriched pathways among downregulated genes in E16.5 cells from Smarcc1/Smarcc2 double knockout mice. **(c)** PPI network of downregulated genes in "chemical synaptic transmission" GO pathway.

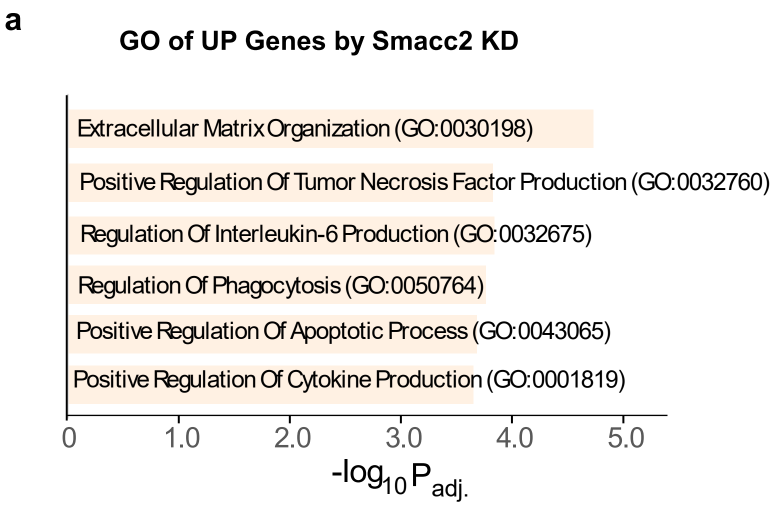


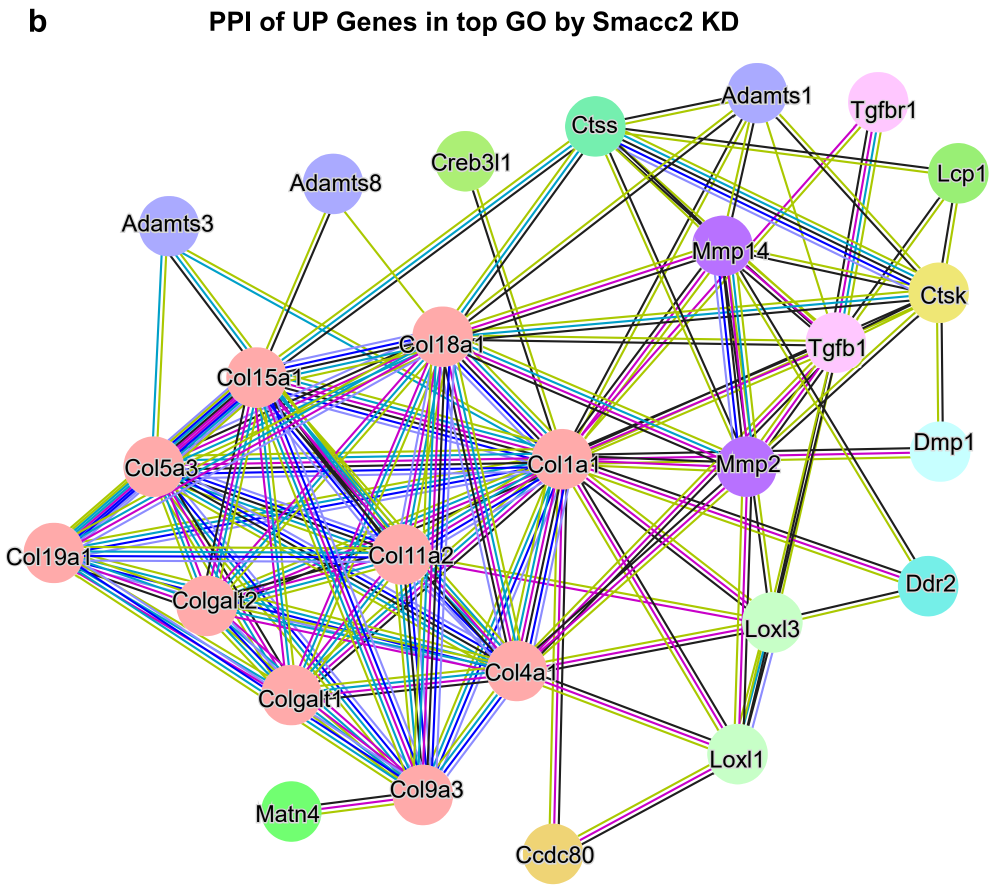


**Figure S3. Smarcc2 knockdown induces the upregulation of genes involved in extracellular matrix and immune response. (a)** Gene Ontology (GO) enriched pathways of upregulated genes in PFC of Smarcc2$^{KD}$ mice, compared to controls. **(b)** PPI network of upregulated genes in top GO category.

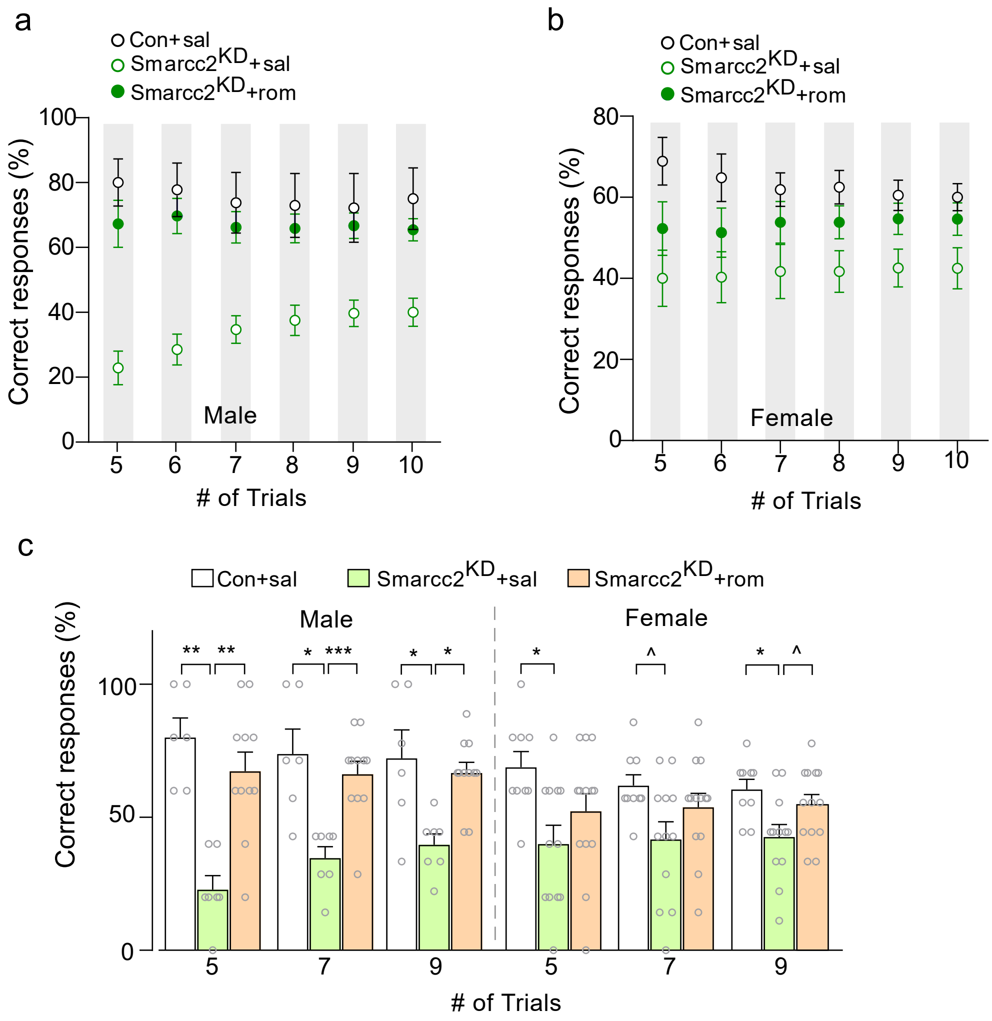


**Figure S4. HDAC inhibitor romidepsin rescues working memory deficits of Smarcc2$^{KD}$ mice of both sexes. (a-c)** Plots showing the percentage of correct responses across trials in T-maze tests of control vs. Smarcc2$^{KD}$ mice of each sex treated with saline or romidepsin. Con+sal: n= 6M, 9F, Smarcc2$^{KD}$+sal: n=7M, 12F, Smarcc2$^{KD}$+rom: n=11M, 13F. All data are shown as mean ± SEM, **p*<0.05, ***p*<0.01, ****p*<0.001. Detailed statistical data are provided in Source Data files.
